# Neutral lipid processing in glia is sexually dimorphic and promotes sleep through diacylglycerol catabolism

**DOI:** 10.1101/2025.09.10.674993

**Authors:** Elana S. Pyfrom, Connor Beveridge, Paula R. Haynes, Vishal A. Kanigicherla, Caitlin E. Randolph, Paula Carvalho Costa, Seble G. Negatu, Sanjay Iyer, Samantha L. Killiany, Zhifeng Yue, Erick N. Astacio, Katherine A. Walker, Kiet N. Luu, Pavel A. Pivarshev, Kellie A. Jurado, Gaurav Chopra, Amita Sehgal

## Abstract

Sleep is thought to have a protective role in clearing toxic waste from the brain, which may include processing of damaged lipids. We recently showed that blocking endocytosis in glia increases sleep and report here that this block is associated with an increase in peroxidized lipids and glial lipid droplet accumulation, raising the possibility that accumulation of these lipids increases the need to sleep. Sleep gain induced by blocking glial transport is exaggerated by knockout of the lipid droplet coat protein, Lipid Storage Droplet 2 (*Lsd2*), suggesting that sleep-promoting lipids are not contained in lipid droplets. To identify lipids regulated by sleep state, we performed global, targeted lipidomics analysis on *Drosophila* neurons and glia, screening nearly 3,000 lipids across 11 major classes. This revealed that sex influences lipid composition in both cell types and lipid homeostasis following extended wakefulness. Female neurons and glia are enriched in ultra-long chain fatty acids, triacylglycerols, and diacylglycerols, with glial diacylglycerol enrichment correlating with elevated sleep need. Based on manipulations of neutral lipid metabolic pathways, we propose that monoacylglycerols, products of glial diacylglycerol catabolism, promote sleep.

## Introduction

Sleep is a conserved behavior that serves critical functions and is present even in simple animals with neural cells but no central nervous system [1]. Metabolism and brain energetics are increasingly linked to sleep and wake dynamics, but the pathways and molecules involved remain largely unknown [2]. In particular, the role of lipids in sleep regulation is underexplored despite the brain having the highest lipid content of any organ after adipose tissue itself [3]. With more neural cells or with centralized organization in the central nervous system, other cell types, primarily glia, may exist to assist neurons in supporting metabolic activity, including lipid homeostasis [4, 5, 6].

A few prior studies support a connection between sleep and lipids. Gerstner et al. showed that fatty acid binding protein (Fabp) [7], which uptakes fatty acids (FAs) into cells and transports them [7], contributes to sleep regulation across species; in mammals, sleep is modulated by glial *Fabp7* [8, 9, 10]. Enzymes that activate FAs increase in expression following sleep loss, and FA-derived lipids may regulate sleep [11, 12, 13, 14]. In *Drosophila*, dietary supplementation of FAs in flies with mitochondrial dysfunction or of acylcarnitines, carnitine-conjugated FAs, increases sleep [15, 6, 16]. A restorative effect of FAs on sleep quality was also demonstrated in a *Drosophila* model of Alzheimer’s disease [17]. Acute sleep loss from mechanical sleep deprivation affects subsequent sleep and memory, and these effects are modulated by the storage of FAs in lipid storage organelles, lipid droplets (LDs) [18]. LDs in glia are themselves regulated by sleep via sleep and apolipoprotein-dependent transfer of lipids from neurons to glia [6]. Thus, localization and storage of lipids shift during wake and sleep, potentially regulating these different behavioral states. In general, we lack understanding of the specific lipids regulated by sleep:wake cycles across different cell types, or the extent to which they contribute to sleep regulation.

We recently reported that blocking endocytosis in *Drosophila* glia, through expression of a dominant-negative *dynamin* allele, shibire, increases sleep [19]. As this manipulation could block the apolipoprotein-dependent lipid transport between neurons and glia described above [6], we asked whether impairments in lipid trafficking account for the increased sleep. Surprisingly, we found that glial endocytosis is not required to accumulate glial LDs as LDs were elevated in brains with the endocytosis block. However, increases in glial LDs are not always associated with increased sleep [6], although specific lipids may be. Using comprehensive metabolic profiling of lipids in neurons and glia across different sleep and wake states, we identify sexually dimorphic regulation of brain lipids, which may underlie differences in sleep between males and females. Finally, we identify a sleep-regulating monoacylglycerol transferase, which we refer to as *tuntid*– the Jamaican Patois word meaning “groggy”. We report a mechanism that promotes sleep by reducing synthesis of the neutral storage lipid, diacylglycerol, from monoacylglycerol by *tuntid* in glia.

## Results

### Perturbing glial transport increases accumulation of brain lipid droplets

We showed previously that blocking endocytosis in glia, using a dominant-negative *dynamin*, increases sleep as well as lipids in the *Drosophila* head [19, 16]. Based on other findings that sleep regulates the transfer of lipids from neurons to glia, where they accumulate as LDs [6], we asked whether endocytosis was required for lipid transfer or LD accumulation in glia. Thus, we measured LD accumulation in female brains with a block in glial endocytosis, produced by Repo-Gal4-driven expression of the dominant-negative *dynamin* allele, *shibire^ts1^*(*shi^ts1^*). Measurements were made at times of high (ZT14) and low (ZT2) glial LD accumulation [6]. We found that pan-glial expression at room temperature leads to increased count, area, and size of LDs at ZT14 compared to genetic controls (Figures 1A-C, 1E). At ZT2, the size of glial LDs as well as brain malondialdehyde levels, marking lipid peroxidation, were increased compared to controls (Figures 1C-D, 1F).

**Figure 1:**
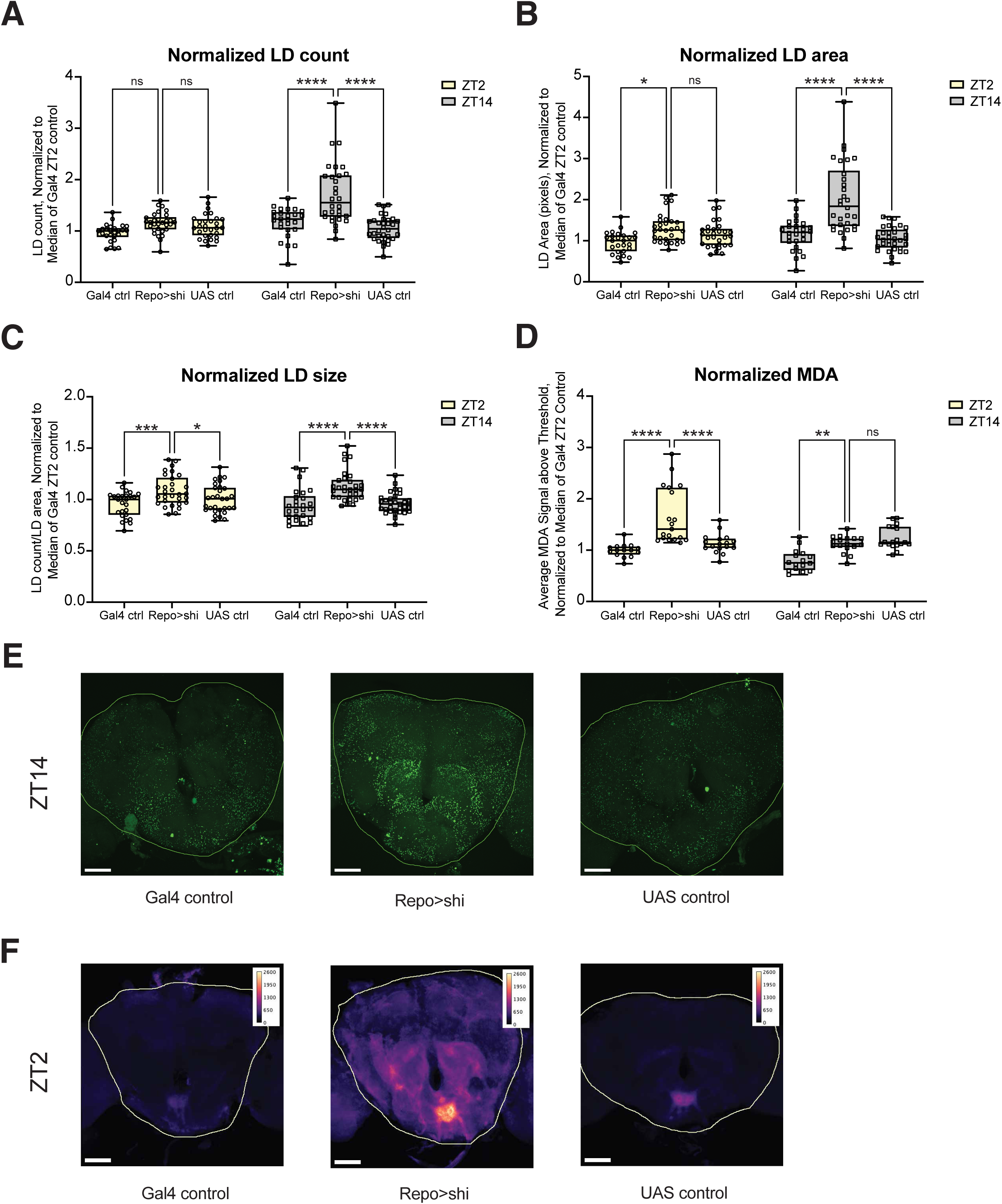
Blocking pan-glial transport increases LD accumulation and malondialdehyde levels. Flies with a pan-glial transport block (Repo-Gal4>20X-UAS-TTS-shi^TS^-p10) and their Gal4 (Repo-Gal4>iso) and UAS (iso>20X-UAS-TTS-shi^TS^-p10) controls were dissected at ZT2 and ZT14 with Bodipy 493/503 brain lipid droplet staining. Normalized LD A) count, B) area, and C) size and D) malondialdehyde staining are shown for an individual fly’s central brain at ZT2 (yellow background) and ZT14 (grey background). E) Maximum projections of representative brains dissected at ZT14 displaying Bodipy 493/503 with a 50 micron scale bar. Central brain is outlined for similar ROI used in analysis. F) Summed projections of representative brains from confocal imaging of anti-MDA antibody staining with 50 micron scale bar. Images are false colored with magma LUT, and central brain is outlined for similar ROI used in analysis. In A-D, all data points are normalized to Gal4 ZT2 median value from its respective replicate. A-C use n = 3 replicates, and all individual brains from all replicates are displayed as data points. D has n = 2 replicates, and all individual brains from all replicates are displayed as data points. The values used for normalization for each replicate are the following: Count (A): 8903.5, 14856, 16074; Area (B): 363039, 460646, 829499; Size (C): 42.2, 31.7, 53.6; and MDA (D): 197.9, 217.1. Two-way analysis of variance (ANOVA) with within-ZT Dunnett’s post hoc test. For all data shown *p≤0.05, ** p≤0.01, *** p≤ 0.001, and **** p≤0.0001, while p≥0.05 is not significant (ns).

Although these findings are consistent with increased acylcarnitine accumulation when glial transport is blocked [16], they are nonetheless surprising in suggesting that functional endocytosis is not required for lipid transfer from neurons to glia.

### LD storage is decreased with reduced exocytosis from cortex glia

Though the *shi^ts1^* allele is used as a tool to block endocytosis, the allele perturbs all functions of dynamin, including exocytosis, and thus may be regarded more as a tool to broadly block vesicular transport [20]. LDs can accumulate when endocytosis is blocked, but their higher level in flies with a glial vesicular transport block (Repo>shi flies) may reflect a failure of exocytosis. We previously found that ecdysone feeding promotes sleep by mobilizing lipids stored in cortex glial LDs and that cortex glia are the major site of sleep-dependent LD accumulation, indicating an important role for these glia [21, 6]. Thus, we knocked down *exo70*, which is required for exocytosis [22], in cortex glia and dissected brains at ZT14, which is when we observe an increase in sleep above control flies with cortex glia-specific *exo70* knockdown (Figure 2A). Measurements of LDs and malondialdehyde levels at ZT14 showed that LD size was significantly decreased compared to controls (Figures 2B-C), while no changes were observed in LD number, area, nor brain malondialdehyde levels (Supp. Figures 1A-C). These data suggest that LD accumulation does not drive sleep need, although the processing of neutral lipids, reflected in reduced size of LDs, may do so. This was supported by assays of flies that express *shi^ts1^* in blood brain barrier (BBB) glia and show increased sleep (Supp. Figure 1D) [19]. We found no changes in LD metrics nor malondialdehyde levels with BBB glia-specific transport block, further disrupting the correlation between LD accumulation and sleep (Supp. Figures 1E-H) [6].

**Figure 2:**
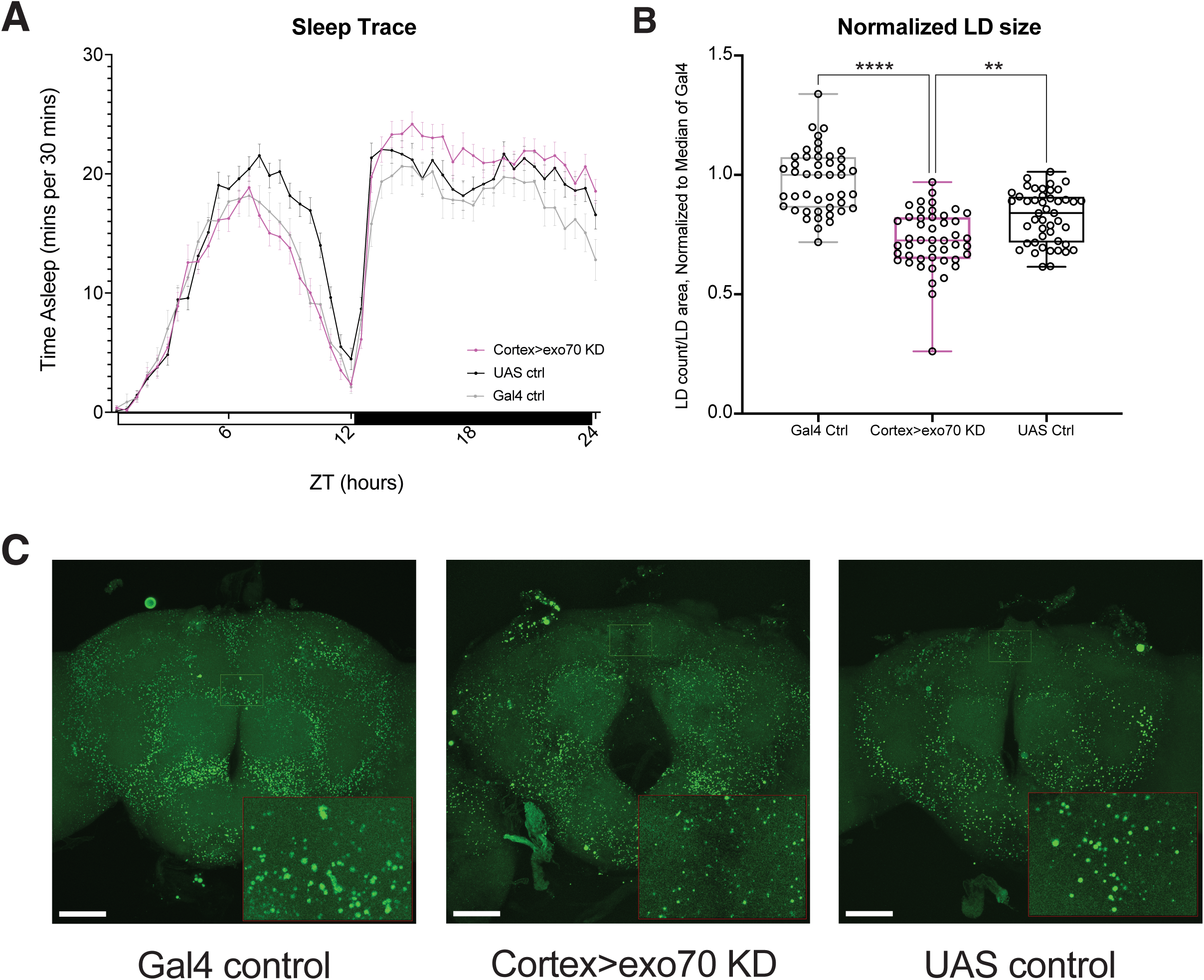
LD size is decreased at ZT14 with reduced exocytosis in cortex glia. Flies with a cortex glia-specific knockdown of exocytosis (iso/y^1^w*; exo70 RNAi/+; cortex-Gal4/+) were assayed for sleep and LDs for comparison to the Gal4 (iso/y^1^w;30B insertion control/+; cortex-Gal4/+) and UAS (iso/y^1^w*;exo70 RNAi/+) controls. A) Sleep trace from females in nutrient-complete, multibeam tubes. B) In brains dissected at ZT14, LD size is decreased in the experimental flies. C) Maximum projections of representative, dissected brains with a 50 micron scale bar. Small green rectangle is drawn around an ROI dorsal to the esophageal foramen. The image is enlarged 4.0 times and found in the red rectangle at the bottom right corner to show differences in LD size. A has n = 1 replicate. B has n = 3 replicates; Kruskal-Wallis with Dunn’s post hoc test, uncorrected. The data points are normalized to Gal4 median value from its respective replicate, and all individual brains from all replicates are displayed. The values used for normalization for each replicate are the following: 31.2, 54.8, 47.6. For all data shown *p≤0.05, ** p≤0.01, *** p≤ 0.001, and **** p≤0.0001, while p≥0.05 is not significant (ns).

### Blocking LD storage exacerbates sleep gains from blocked glial transport

To confirm that LD accumulation does not drive sleep in Repo>shi flies, we manipulated triacylglycerols (TGs), the primary neutral storage lipid found in LDs by introducing a *brummer* loss-of-function mutation *(bmm^1^)* into this background. *bmm* is the *Drosophila* homolog of *adipose triglyceride lipase, atgl*, and loss of *bmm* reduces the breakdown of TGs and leads to more LD storage [23]. We first tested *bmm^1^* alone for effects on sleep. *bmm^1^* mutations were previously reported to have no effect on baseline sleep but increase the homeostatic sleep response following sleep deprivation [18]. Using a sleep assay that monitors sleep and wake states with higher resolution (multibeam *Drosophila* activity monitors) and our standard sucrose-based diet, we find similar results for baseline sleep in females (Figure 3A), although *bmm^1^* males have markedly reduced total sleep (Figure 3B). To mimic the previous sleep study with *bmm^1^* mutants, we performed sleep assays in single beam monitors on nutrient-complete food [18]. While females show no change in sleep as previously reported (Supp. Figure 2A), *bmm^1^* males still show significant sleep loss compared to control males (Supp. Figure 2B). This suggests that previously reported sexual dimorphism in Bmm activity has sex-specific consequences for sleep [24, 25].

**Figure 3:**
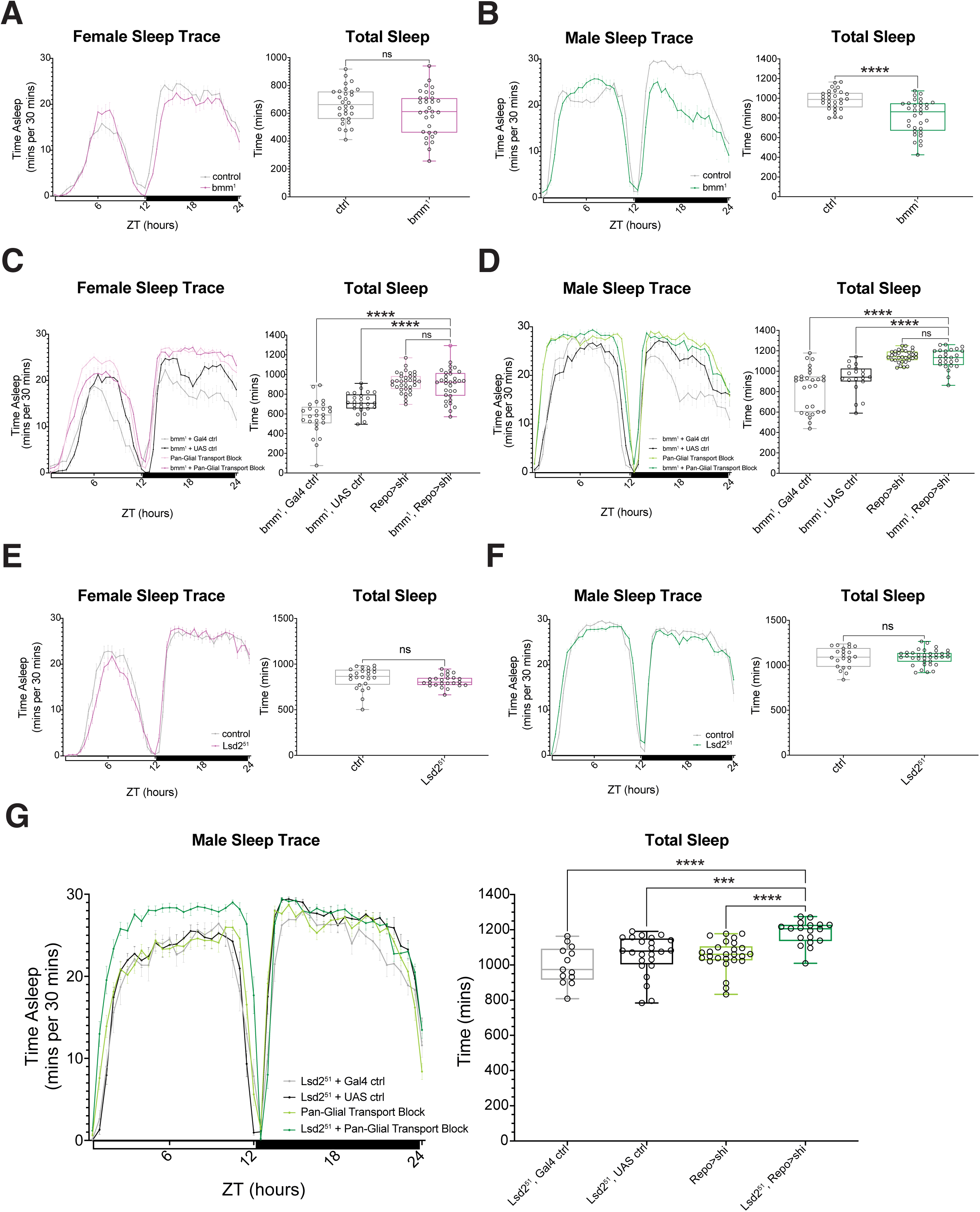
Simultaneously blocking LD formation and glial transport increases sleep. Averaged multibeam sleep traces and total sleep on sucrose food for *bmm^1^* mutants (w’/w*(iso);;bmm^1^, Repo-Gal4/bmm^1^) and controls (w’/w’(iso);;Repo-Gal4/+). A) No change in sleep is observed in females and B) males have decreased sleep. C) Females and D) males with both pan-glial transport block and bmm^1^ homozygous mutation (*w’;;bmm^1^, Repo-Gal4>20X-UAS-TTS-shi^TS^-p10/bmm^1^*) was compared to Gal4 (*w’;;bmm^1^, Repo-Gal4/bmm^1^*), UAS (*w’;;bmm^1^, 20X-UAS-TTS-shi^TS^-p10/bmm^1^*), and mutant (*w’;;Repo-Gal4/20X-UAS-TTS-shi^TS^-p10*) controls. Averaged multibeam sleep traces and total sleep show the experimental groups for both sexes sleep similar amounts as flies with a pan-glial transport block. E) Females and F) males with *Lsd2^51^* homo/hemizygous mutations (*y^1^, Lsd2^51^;; Repo-Gal4/+*) averaged multibeam sleep traces and total sleep compared to controls (*y^1^;; Repo-Gal4/+*). G) Averaged multibeam sleep traces and total sleep for males with both pan-glial transport block and Lsd2^51^ hemizygous mutation (*y^1^, Lsd2^51^/Y;; Repo-Gal4>20X-UAS-TTS-shi^TS^-p*10) was compared to Gal4 (*y^1^, Lsd2^51^/Y;; Repo-Gal4/+*), UAS (*y^1^, Lsd2^51^/Y;; 20X-UAS-TTS-shi^TS^-p10/+*), and mutant (*y^1^/Y;; Repo-Gal4/20X-UAS-TTS-shi^TS^-p10*) controls, showing experimental group’s sleep is increased above all controls. A and B have n = 3 replicates; Mann-Whitney test (two-tailed). C and D have n = 3 replicates; Kruskal-Wallis test with Dunn’s post hoc test, uncorrected. E has n = 3 replicates; Mann-Whitney test (two-tailed). F has n = 6 replicates; Mann-Whitney test (two-tailed). G has n = 3 replicates, where two replicates have 10-12 flies for the Lsd-2 Gal4 control; Kruskal-Wallis test with Dunn’s post hoc test, uncorrected. All figures display results from one representative replicate with data points representing individual flies. For all data shown *p≤0.05, ** p≤0.01, *** p≤0.001, and **** p≤0.0001, while p ≥ 0.05 is not significant (ns).

We next coupled the *bmm^1^* mutation with the pan-glial block in transport, predicting that increased TGs in LDs would not further increase sleep in Repo>shi flies. *bmm^1^* mutation did not further increase nor suppress the sleep phenotype of Repo>shi flies (Figures 3C-D). Rather, flies with *bmm^1^* mutation and glial transport block slept similarly as flies with Repo>shi alone. This was even true in males, despite the reduced sleep of *bmm^1^* males at baseline (Figure 2B).

Finally, we blocked LD storage by mutating *Lipid storage droplet-2 (Lsd2)*, the *Drosophila* homolog of *Perilipin-2*. *Lsd2* localizes to the endoplasmic reticulum to stabilize the formation and growth of LDs, and *Lsd2* can also translocate to LDs itself to regulate Bmm lipase activity and ultimately LD size [26]. *Lsd2^51^* mutants were reported to have reduced TG levels, normal baseline sleep, and reduced sleep after sleep deprivation [27, 18]. We found that *Lsd2^51^* mutant females trend towards decreased sleep, but there was no robust, significant change, which is consistent with the previous study (Figure 3E) [18]. *Lsd2^51^* mutant males display inconsistent sleep, showing either no change or strikingly decreased sleep (Figure 3F shows an example of unaffected sleep). Although combining the X-linked *Lsd2^51^* mutation with the *shi^ts1^* allele proved technically challenging in females, addition of a *Lsd2^51^* mutation to Repo>shi males produced a synergistic effect by dramatically increasing sleep (Figure 3G). These data support the idea that LDs themselves do not promote sleep, but rather that lipids affect sleep prior to storage in LDs. We suggest that neutral lipid processing promotes sleep.

### Sex is a primary regulator of Drosophila brain lipid composition

To investigate how the glial lipidome changes with sleep need, we dissected brains from alert flies (ZT2) and two conditions with high sleep need: “sleepy” flies, which are collected after the wake period (ZT14), and sleep-deprived (SD) flies– subjected to 10-hour mechanical sleep deprivation from ZT16 to dissection at ZT2 (Figure 4A). Flies expressed mCD8-GFP in either neurons (Nsyb-Gal4>UAS-mCD8-GFP) or glia (Repo-Gal4>UAS-mCD8-GFP) to label cells for Fluorescence-activated cell sorting (FACS) (Figure 4A, Supp. Figure 3A-B). For each replicate, 100,000 GFP-positive cells were collected per condition, stratified by sleep state, cell type, and sex (Figure 4A, Supp. Figure 3C-D).

**Figure 4:**
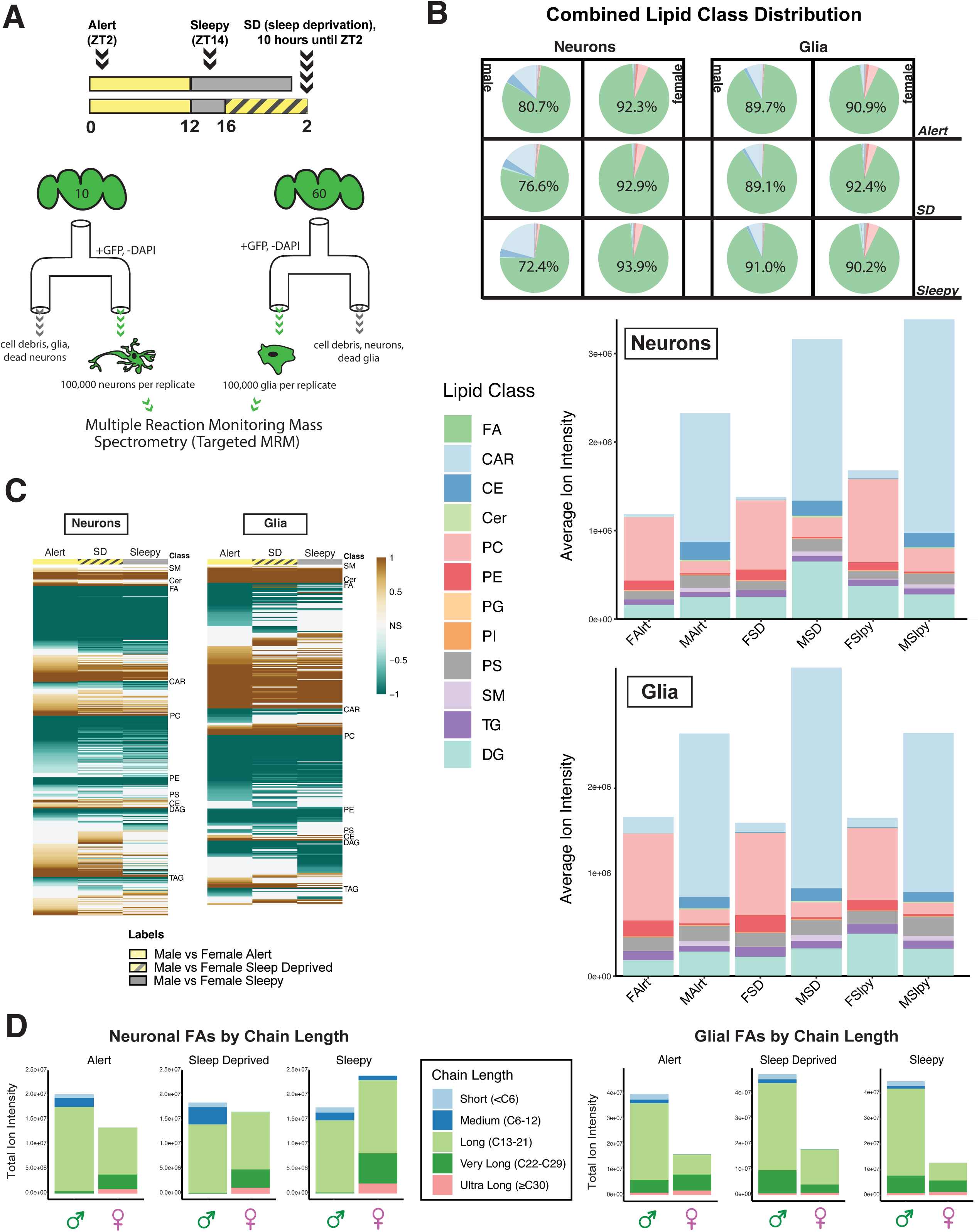
Sex is a primary source of variance in *Drosophila* brain lipid composition. A) Simplified schematic of protocol for *Drosophila* brain cell-sorted lipidomics. 10 brains per sleep condition expressing +/(w’ or Y);;Nsyb-Gal4>20X-UAS-mCD8-GFP and 60 brains per sleep condition expressing +/(w’ or Y) ;;Repo-Gal4>20X-UAS-mCD8-GFP were dissected. Approximately 100,000 neurons and glia per condition, per replicate were collected through FACS (GFP-positive, DAPI-negative) for targeted multiple reaction monitoring mass spectrometry. The three conditions include brains dissected at ZT2 (Alert), ZT14 (Sleepy), and ZT2 following 10 hours of sleep deprivation (SD). Flies were entrained to a 12 hour light (yellow rectangle):12 hour dark (grey rectangle) cycle, with 10 hour sleep deprivation (for 8 hours during dark and 2 hours during light cycle) from ZT16 to ZT2 (yellow and grey stripes). B) Proportions of lipids detected by targeted multiple reaction monitoring mass spectrometry (MRM) are separated by class in two ways: (Above) The pie chart displays the average intensity of each lipid class, revealing that FAs exhibited the highest relative signal. Neuronal samples are displayed on the left and glial samples on the right. Columns alternate by sex, with males in the first and third columns and females in the second and fourth columns. Rows represent different sleep states: alert samples in the first row, sleep-deprived samples in the second row, and sleepy samples in the third row. The relative percentage of FAs (green) is indicated within each respective box. (Below) A bar plot displays the average intensity of lipid classes per condition, excluding fatty acids (FAs). The conditions are abbreviated as follows: FAlrt (Female Alert), MAlrt (Male Alert), FSD (Female Sleep Deprived), MSD (Male Sleep Deprived), FSlpy (Female Sleepy), and MSlpy (Male Sleepy). Neuronal samples are shown above glial samples. Lipid classes in panel B include: CAR (Acylcarnitines, light blue), CE (Cholesterol esters, dark blue), Cer (Ceramides, light green), PC (Phosphatidylcholines, coral), PE (Phosphatidylethanolamines, red), PG (Phosphatidylglycerols, light orange), PI (Phosphatidylinositols, orange), PS (Phosphatidylserines, grey), SM (Sphingomyelins, lavender), TG (Triacylglycerols, dark purple) and DG (Diacylglycerols, turquoise). C) The Log Fold Change (LogFC) heatmap displays differentially expressed lipid species comparing males versus females across three sleep conditions: alert (ZT2), sleep-deprived (SD), and sleepy (ZT14) flies. Brown coloring indicates upregulation (LogFC is ≥ 1) in males relative to females, while teal indicates downregulation (LogFC ≤ -1) in males relative to females. Non-significant lipids are white. D) Total FA intensity detected for neuronal and glial fatty acids color-coded by chain length. Short chain FAs have lengths less than 6 carbons (light blue), medium chain range from 6-12 carbons (dark blue), long range from 13-21 carbons (light green), very-long from 22-29 carbons (dark green), and ultra-long chain are FAs with 30 or greater carbon chains (pink). Each cell type and sleep condition subfigure depicts two stacked bars with males being on the left (green circle and arrow underneath) and females being on the right (pink circle with intersecting lines underneath).

Targeted multiple reaction monitoring (MRM) profiling screened for over 3,000 lipid species, with assignments guided by characteristic fragmentation patterns and class-specific ionization properties (Figure 4A). To visualize lipid composition, we normalized signal intensities for each lipid class by dividing the total signal by the number of monitored MRMs (Figure 4B). Overall separation by sex was clear: principal component analysis (PCA) indicated that sex explained more variance than either sleep state or cell type, although PC1 and PC2 together accounted for only ∼30% of total variance (Supp. Figure 3E). This strong sex effect was reinforced by a heatmap of differentially expressed (FDR < 0.1) lipids, which revealed striking male–female separation across all sleep states and cell types (Figure 4C). Log fold change (LogFC) heatmaps revealed that males exhibited broad upregulation of lipid features across sleep states (Figure 4C), including increased FA abundance in both neurons and glia, with male-biased enrichment especially evident in glia (Figure 4D). Although females displayed a greater percentage of FA signal in both neurons and glia compared to males, this is likely due to the high abundance of acylcarnitines detected in males (Figure 4B).

Within this overarching sex difference, conditions of high sleep need (sleepy and SD) were associated with a decrease in neuronal long chain FAs (LCFAs, 13-21 carbons or C13-21) in males and an increase in females compared to the alert condition; a male-specific increase in glial LCFAs was also detected (Figure 4D). Very long chain FAs (VLCFAs) were strongly enriched in females, particularly ultra-long-chain FAs (ULCFAs,

≥C30), a subset of VLCFAs [28, 29]. Neuronal VLCFAs (C22-29) and ULCFAs were scarcely detected in males but readily detected in females, with levels increasing with high sleep need (Figure 4D). In females, glial VLCFAs were relatively higher as a proportion of total FAs, but they decreased with sleep need while increasing in males (Figure 4D). Despite an overall lower FA signal in glia from females, ULCFAs remained more abundant in females than in males across sleep states (Figures 4D). Because FA profiling relied on a pseudo-MRM approach in which identical precursor/product ion pairs were monitored, overlap with other anionic lipids is possible; thus, FA annotations should be considered provisional pending orthogonal validation.

### Sleep-regulated brain lipids are sexually dimorphic

We next examined how lipid profiles change with sleep state within each sex by comparing alert flies to high sleep need groups (sleepy and sleep-deprived). Differential expression analysis (FDR < 0.1) revealed widespread changes in both neurons and glia with distinct, sex-specific patterns (Supp. Figure 4A). In females, several TG species were upregulated in both neurons and glia under conditions of high sleep need (Table 1). In males, more changes in lipid features were observed in neurons, but DGs were consistently elevated across all sleep need conditions relative to alert (Table 1). Glial DG accumulation was especially pronounced in females (Supp. Figure 4B), where DGs comprised a third of the most strongly regulated species (Table 1), supporting a sex-modulated role for DG metabolism in promoting sleep. Glycerophosphocholines (PCs), known precursors of DGs [30], were elevated in neurons of both sexes with sleep need but decreased in female glia (Figure 4B), suggesting potential crosstalk between sex and sleep-dependent lipid regulation.

**Table 1:**
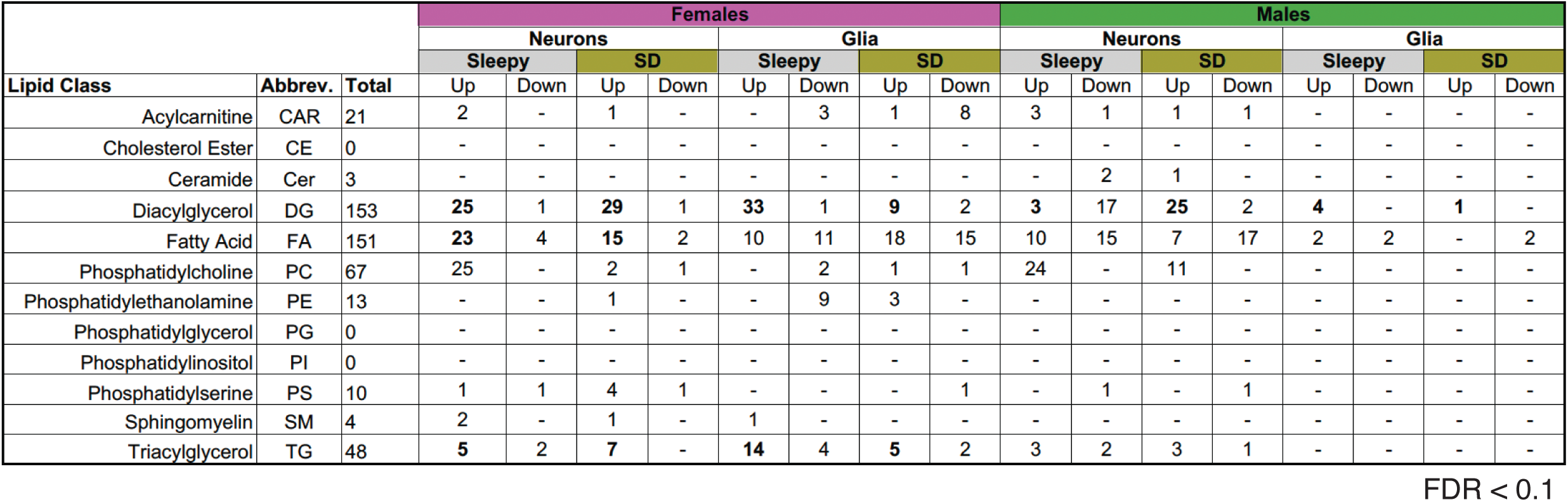
Lipid species hits by class compared to alert (ZT2) condition. Values represent total number of lipid species within each lipid class that has a False Discovery Rate (FDR) < 0.1. Up and Down refer to whether the lipid species is increased (up) or decreased (down) in the condition with increased sleep need when compared with alert (ZT2) flies with the same sex and cell type.

FA differential regulation also varied with sleep state, though patterns were less consistent across sex, cell type, or condition besides an abundance increase in FA abundance in female neurons with sleep need (Table 1). Several of the top lipid features were annotated as C14 and C15 FAs (Supp. Table 1). Such chain lengths are biologically relevant because they can participate in protein acylation through myristoylation (C14) and farnesylation (C15) [31, 32]. Features annotated as C14 FAs were enriched in alert males, whereas C15 features were elevated in females and enriched with high sleep need in both sexes (Supp. Table 1, Supp. Figure 5A).

To explore whether protein acylation changes with sleep need, we examined myristoylation and farnesylation with commercial antibodies. Only the farnesylation antibody produced a specific signal in flies (Supp. Figure 5B) [33]. Imaging of central brains revealed a trending decrease of farnesylation in sleepy males and a significant reduction following sleep deprivation (Supp. Figure 5C-D). In contrast, females showed a trend toward increased farnesylation with sleep loss, marked by small, bright punctae observed almost exclusively in sleep-deprived females (Supp. Figure 5E). These results highlight sex-specific regulation of farnesylation and support the interpretation that FAs linked to acylation, particularly C15, are modulated by both sex and sleep.

### Analysis of diacylglycerol metabolism negates a sleep-promoting role for diacylglycerol

Our lipidomic data indicated glial DG accumulation following prolonged wake, particularly in females (Figures 3A, 3C, and Supp. Figure 4B; Table 1). These findings implicated DGs in sleep regulation. To test whether DGs themselves or their metabolic products promote sleep, we manipulated DG metabolic pathways in females.

DGs are reduced in the retina when *roundabout (rbo)*, a lipase and vesicular transport-related enzyme that scaffolds phosphatidylinositols, is knocked down [34, 35, 36]. Adult-specific, glial knockdown of *rbo* reduced female sleep, consistent with a role for DG accumulation in promoting sleep (Figure 5A). To increase DG levels, we knocked down *inaE*, the only *Drosophila* DG lipase with homology to mammalian DG Lipase alpha and beta [37, 30]. Unexpectedly, adult-specific knockdown of *inaE* in glia reduced female sleep (Figure 5B). *Drosophila* have a genomic cluster where three paralogous genes, *CG1941*/*tuntid*, *CG1942*/*bishu-1*, and *CG1S4C*/*bishu-2*, share homology with the mammalian Mogat2 and Dgat2 family, which synthesize DGs from monoacylglycerols (MG)s and TGs from DGs, respectively [24, 38, 39]. Knockdown of *bishu-2* in S2 cells results in DG accumulation [40], so we knocked it down in female glia as another method of increasing glial DGs. Glial *bishu-2* knockdown results in no change in sleep (Figure 5C), which supports the previous finding that DGs are not the sleep-promoting lipid in glia.

**Figure 5:**
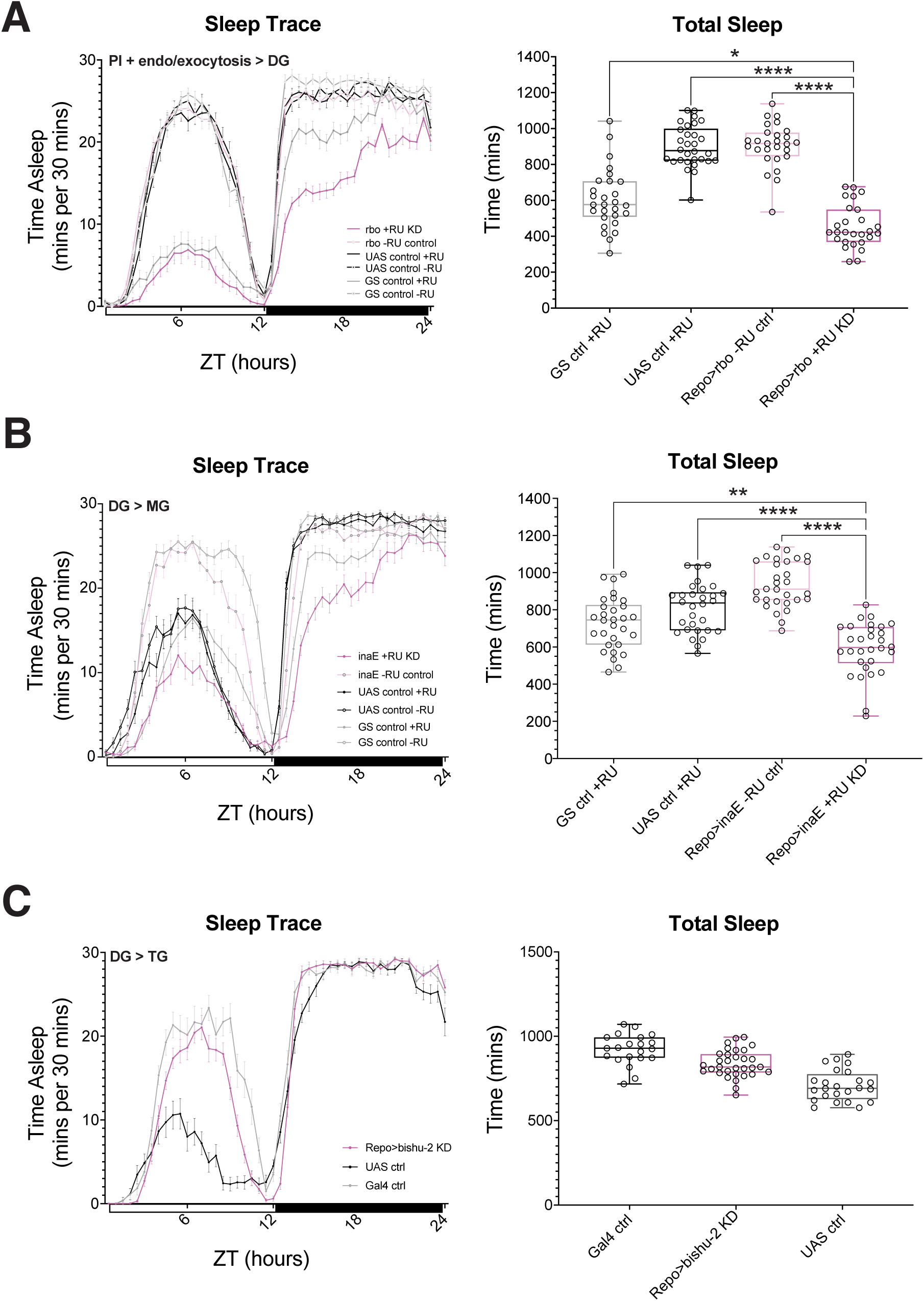
DG accumulation in glia does not promote female sleep. Averaged multibeam sleep traces and total sleep for females on sucrose-based food. In A) and B), 100 uM RU-486 (+RU) and ethanol (-RU) control was fed to females with glial conditional knockdown of A) *rbo* and B) *inaE* using the GeneSwitch (GS) system. Total sleep analysis compares the experimental group on +RU food with genetic controls (GS and UAS controls) on +RU food and the experimental (Repo-GS>UAS-RNAi) without RU (-RU) to determine adult-specific effects. A) Adult-specific, glial knockdown of *rbo* (wCS/w^1118^; P{GD14013}v47751/+; Repo-GS/+, +RU) has decreased total sleep compared to GS control (w’CS;;Repo-GS/+, +RU), UAS control (w’CS/ w1118; P{GD14013}v47751/+, +RU), and drug control (w’CS/w1118; P{GD14013}v47751/+; Repo-GS/+, -RU). B) Adult-specific, glial knockdown of *inaE* (w’CS/y^1^, sc*, v^1^, sev^21^; P{TRiP.HMC05758)attP40/+ ; Repo-GS/+, +RU) has decreased sleep total compared to GS control (w’CS/ y^1^, v^1^; P{Msp300}attP40/+; Repo-GS/+, +RU), UAS control (w’CS/y^1^, sc*, v^1^, sev^21^; P{TRiP.HMC05758)attP40/+, +RU), and -RU control (w’CS/y^1^, sc*, v^1^, sev^21^; P{TRiP.HMC05758)attP40/+ ; Repo-GS/+, -RU). C) Pan-glial knockdown of *bishu-2/CG1S4C* (w’CS/y^1^w*; P{KK112067}VIE-260B/+; Repo-Gal4/+) shows no sleep change compared to Gal4 (w’CS/y^1^w*; 30B insertion control/+;Repo-Gal4/+) and UAS (w’CS/y^1^w*; P{KK112067}VIE-260B/+;+) controls. All figures display results from one representative replicate with data points representing individual flies. RU486 experiment shows sleep traces for additional -RU controls. A has n = 2 replicates; Kruskal-Wallis test with Dunn’s post-hoc test, uncorrected. B has n = 3 replicates; Kruskal-Wallis test with Dunn’s post-hoc test, uncorrected. C has n = 1 replicate; Kruskal-Wallis test with Dunn’s post-hoc test, uncorrected. For all data shown *p≤0.05, ** p≤0.01, *** p≤0.001, and **** p≤0.0001, while p ≥ 0.05 is not significant (ns).

### Monoacylglycerol accumulation in glia promotes sleep

Because we find that DGs accumulate in glia with sleep need and that glial knockdown of the DG lipase, *inaE* leads to sleep loss, we investigated whether products of DG catabolism promote sleep. *inaE* hydrolyzes DGs into monoacylglycerols (MGs), which in *Drosophila* are metabolized by multiple enzyme families, including fly homologs of SERHL2, ABHD12, and ABHD6 [41]. We asked whether MGs are the relevant sleep-promoting species by supplementing the diet with NF1819, an irreversible inhibitor that targets multiple MG lipases [42, 43]. NF1819 supplementation significantly increases sleep in females (Figure 6A), but, in contrast, decreased sleep in males (Supp. Figure 6A).

**Figure 6:**
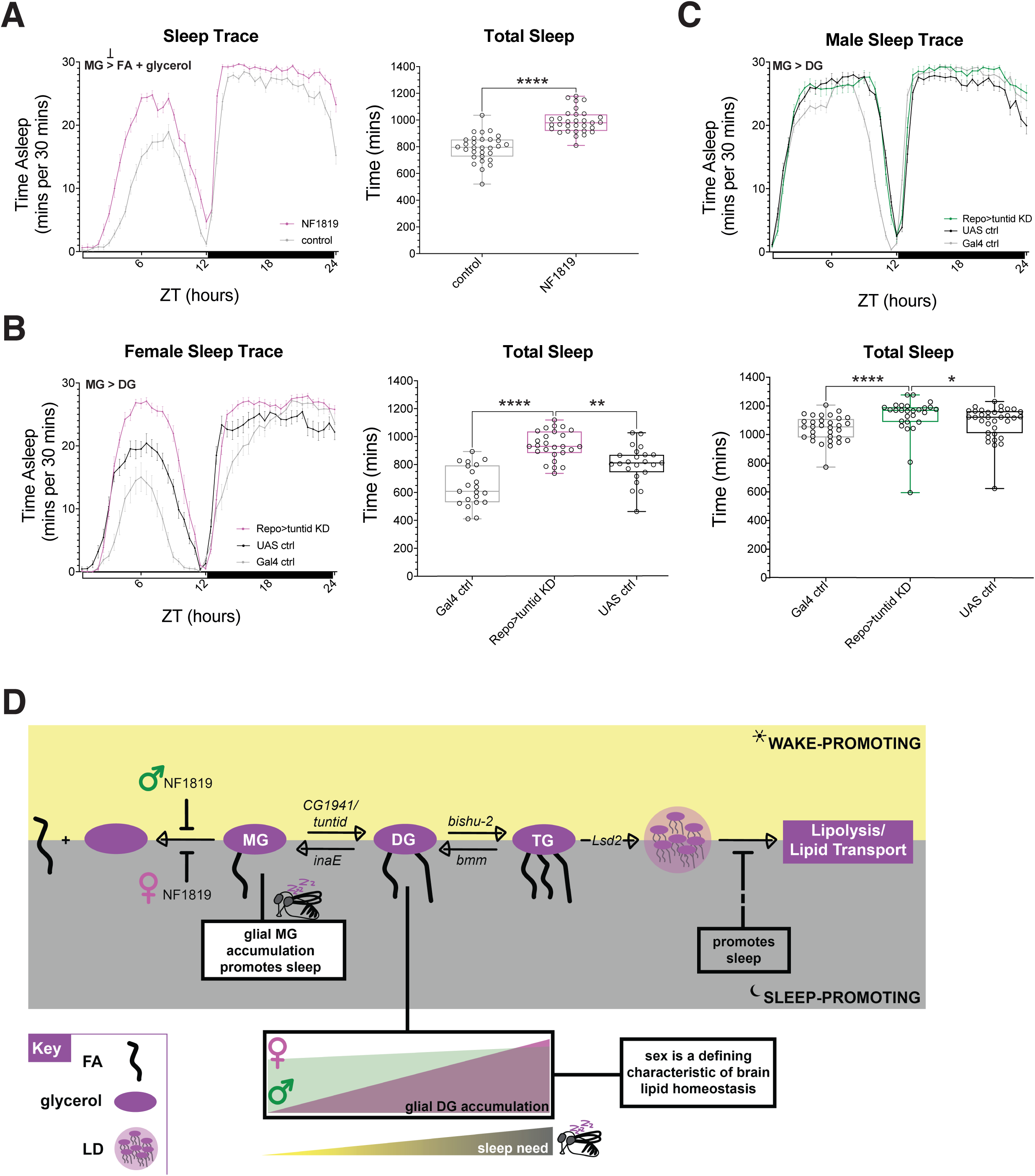
Monoacylglycerol accumulation in glia promotes sleep. Averaged multibeam sleep traces and total sleep for females on sucrose-based food. A) 500 ug/mL NF1819 supplementation increases sleep in Canton S females compared to ethanol control diet. B) and C) Pan-glial knockdown of CG1941 (w’CS;w^1118^, P{GD1853}v3998;; Repo-Gal4/+) increases sleep compared to Gal4 (w’CS;;Repo-Gal4/+) and UAS (w’CS/w^1118^, P{GD1853}v3998) control B) females and C) males. D) Model summarizing findings where sex determines extent of DG accumulation with sleep need, leading to female-specific MG accumulation. Female symbol (pink, circle with intersecting lines below) and male symbol (green, circle with arrow above) denote sex-specific differences in findings, contributing to a model proposing that inhibiting a step(s) upstream of lipolysis and/or lipid transport can promote sleep. Abbreviations include MG (Monoacylglycerol), DG (Diacylglycerol), TG (Triacyglycerol), FA (fatty acid), and LD (lipid droplet). All figures display results from one representative replicate with data points representing individual flies. A has n = 3 replicates; Mann-Whitney test (two-tailed). B has n = 3 replicates; Kruskal-Wallis test with Dunn’s post-hoc test, uncorrected. C has n= 2 replicates; Kruskal-Wallis test with Dunn’s post-hoc test, uncorrected. For all data shown *p≤0.05, ** p≤0.01, *** p≤0.001, and **** p≤0.0001, while p ≥ 0.05 is not significant (ns).

To further test the MG hypothesis, we tested monoacylglycerol transferases (MGATs), enzymes that convert MGs to DGs, for effects on sleep. We knocked down *CG1941*/*tuntid*, which is a homolog of mammalian monoacylglycerol O-acyltransferase, *Mogat2* and whose knockdown strongly suppresses DG production in S2 cells [40]. MGATs are highly expressed in the gut [30], where we validated the RNAi line by qPCR (Supp. Figure 6B). We knocked down *tuntid* in *Drosophila* glia and find that both females (Figure 6B) and males (Figure 6C) have increased sleep, although the effect is not as pronounced in males. *bishu-1* is a paralog of *tuntid*, although its knockdown reduces DGs to a lesser extent than *tuntid* in S2 cells [40]. We knocked down *bishu-1* in glia using two RNAi lines. Glial knockdown (KK construct) of *bishu-1* increases sleep in females and males, but to a lesser extent than *tuntid* (Supp. Figures 6C-D). Knockdown of *bishu-1* with a different RNAi (GD construct) did not increase sleep on sucrose food (Supp. Figures 6E-F) but did so in females on nutrient-compete food (Supp. Figure 6G). Males with glial *bishu-1*-GD knockdown showed unchanged or occasionally decreased sleep (Supp. Figure 6H shows unchanged sleep). Since Tuntid is the most effective at reducing DG levels in S2 cells and also influences the levels of *bishu-1* and *bishu-2* [40, 38], it may be the major MGAT in *Drosophila*. Knockdown of MGATs promotes sleep via MG accumulation in glia, especially in females. This is supported by experiments where MGs are increased through reduced conversion of MGs to DGs (knockdown of *tuntid* and *bishu-1*) and through inhibition of monoacylglycerol lipases (NF1819). Together, these findings demonstrate a sleep-promoting role for MGs and suggest that neutral lipid metabolism underlies at least some sex-specific differences in sleep.

## Discussion

Consideration for how neurons differ from other cell types could provide insight into primary functions of sleep. Neurons’ strikingly complex cell morphologies, uncommon or neuropathological storage of lipids in lipid droplets, reliance on glia for lipid homeostasis, and low capacity for fatty acid breakdown by beta-oxidation indicate that neuronal lipid metabolism and function is regulated differently than in other cell types [3, 44, 4, 45, 6]. We recently identified a neuron-glia lipid metabolic cycle where lipid damage accumulates with prolonged wake, and damage is transferred from neurons to glia by the *Drosophila* homolog of *ApoE*, *Nlaz*. Here, using targeted, brain cell-sorted lipidomics, we find that DG accumulation in the brain is associated with high sleep need. The DGs are likely catabolized to MGs, which we identify as sleep-promoting lipid entities through genetic and pharmacological analysis of neutral lipids in glia. MGs were not included in the targeted lipidomic analysis, but consistent with their derivation from DGs, we find that both species are more associated with sleep/sleep need in females. Notwithstanding the sexually dimorphic lipid composition of female and male brains, we propose MGs as a sleep-promoting lipid, where sleep is modulated by MG to DG conversion by glial *tuntid*.

We investigated LD storage as a primary mechanism within neutral lipid metabolism that promotes sleep, since LD count, area, and size increase with a pan-glial transport block that elevates sleep and sleep need (Figure 1) [19]. Consistent with the results of the pan-glial block, we previously showed that LD metrics in cortex and ensheathing glia increase with prolonged wake, another condition of high sleep need [6]. Reasoning that LD accumulation might drive sleep, we blocked exocytosis from cortex glia and found that while sleep increases, LD size actually decreases, which may be indicative of lipid processing (Figure 2). We also find that malondialdehyde levels, though increased with a pan-glial transport block, did not change with a knockdown of *exo70* (Figure 1D, Supp. Figure 1A). Although we have shown that lipid peroxidation contributes to a sleep-dependent neuron-glia lipid cycle [6], and others have found that lipid peroxidation in neurons can promote sleep [46], the mechanism by which lipids promote sleep through glia may not require peroxidized lipid damage. Rather, with our manipulations, we are likely trapping lipids in cortex glia, requiring their breakdown or processing in these cells. These data support the previous finding that mobilization of lipids in cortex glia promotes sleep [21].

While lipids generally enter glia by endocytic mechanisms mediated by dynamin [45, 47], other modes of transport such as endocytosis not requiring dynamin, like macropinocytosis, or breakdown of lipids into classes that can be taken up by transporters are also possible. Carnitine transporters expressed by BBB glia mediate the promotion of sleep [16, 48], so it is possible that other glia also use transporters, like sleep-modulating Fabp, when endocytosis is blocked [10]. This work supports the idea that neutral lipid metabolism, which may include acylcarnitine metabolism, promotes sleep [49, 16, 48].

Acylcarnitines are generally enriched in males, showing increases with sleep need that are most apparent in male neurons (Figure 4B). While acylcarnitine changes are relatively mild in females, they become more pronounced with high sleep need; however, FAs, DGs, and TGs showed more dramatic alterations (Figure 4B, Table 1, Supp. Figure 4B). Elevated levels of both acylcarnitines and DGs were previously reported in sleep-deprived humans, where a DG species was the most highly accumulating lipid identified [50]. Our study identifies numerous changes in neutral lipid homeostasis by lipidomic analysis.

DG anabolism consists of producing TGs, which are typically stored in LDs [39, 30]. As a means of increasing or reducing lipid storage in LDs, we introduced *bmm^1^* and *Lsd2^51^* mutations, respectively, into the glial transport block paradigm. In this context, loss of *bmm* had no effect (Figure 3C-D), even though it reduced sleep by itself in males (Figure 3B, Supp. Figure 2B), suggesting that sleep promotion by the pan-glial block is epistatic to the decrease in sleep with *bmm^1^*. The *Lsd2^51^* mutation displays a synergistic effect, further promoting sleep in a non-additive manner (Figure 3G). This suggests that sleep is promoted by lipids prior to their storage in LDs. As noted, *bmm^1^* mutants were previously reported to have no change in baseline sleep [18], but it is possible that females were assayed in that case, since we observe male-specific loss of sleep (Figure 3A-B, Supp. Figure 2A-B). Since males have increased Bmm lipase activity [24], they may be more sensitive to loss of *bmm*, which could account for their reduced sleep. Given that Lsd2 can also change subcellular localization to regulate Bmm activity and LD size [26], sexually dimorphic sleep could be due to sex-specific differences in TG breakdown, production, and LD localization of these enzymes [39].

We investigated DGs as the neutral lipids which accumulate to promote sleep. By lipidomic analysis of FACS-sorted neurons and glia, we find an increase in DG signal across all conditions of increased sleep need compared to alert flies (Figure 4B, Supp. Figure 4B). Multiple DG species were differentially expressed in female glia under high sleep need conditions compared to alert samples (Table 1). With sleep need, TG species were also increased in female glia and overall in males (Table 1, Supp. Figure 4B). While we expect increased TGs with prolonged wake because we have reported increased LDs in female glia [6], we note that the primary circulating neutral lipids in *Drosophila* are DGs [51, 52]. Thus, DG accumulation in the brain with high sleep need could also be a mechanism of peripheral tissues communicating a need to sleep through inter-organ lipid signaling with the brain.

Although not pursued here, neuronal DGs are also elevated in all conditions of sleep need, especially in males, despite the presence of more differentially expressed neuronal DG species in male alert conditions than sleepy male neurons (Supp. Figure 4B, Table 1). Neuronal activity is regulated by calcium signaling, which is closely linked to DG metabolism [30]. Another enzyme, Dgat1, integrates calcium signals to produce TGs for LD storage from dietary lipids, indicating other mechanisms by which DG metabolism could contribute to sleep and wake state [30, 53]. Elevated DG signal in both neurons and glia with increased sleep need could also be due to DG transport between brain cell types (Supp. Figure 4B).

In neurons, increased DG signal and high sleep need is associated with increased PC signal, while PC levels decrease in glia with sleep need (Figure 4B, Table 1). PC metabolism closely links DGs with sphingolipid metabolism, and here, we detect low signals of ceramides and sphingomyelins (SMs) in males (Figure 4B). We find sex-specific expression of SM (Figure 4C), which should be validated with more in depth mass spectrometry methods [54]; notably, ceramides and other sphingolipids were shown to be enriched in *Drosophila* glia with prolonged wakefulness [55]. DG metabolism is also closely linked with other phospholipid metabolism, including phosphatidylethanolamines (PEs) and phosphatidic acid (PA). The conserved DG kinase, Dgk, converts DGs to PA by phosphorylation and was recently shown to act in *Caenorhabditis elegans* neurons to mediate sleep [56]. Our lipidomic data show time-of-day differences for PEs and sleep need-dependent differences for PCs, which may suggest DG metabolism as a lipid pathway that allows for crosstalk between brain cells and between circadian and sleep regulation of lipid homeostasis. We find that conditions of sleep, sex, and cell type also interact to maintain lipid homeostasis. FAs commonly used for acylation are often the most changed lipids, though the pattern is not always shared by sex (Supp. Table 1, Supp. Figure 5A). Myristoylation is a type of reversible protein acylation that may change with sleep and wake state in a sex-specific manner, since FAs with 14-carbon chain length are some of the most changed lipid species in alert males (Supp. Table 1). Myristoylation changes the biophysical properties of a protein, altering its membrane binding capabilities, which may affect subcellular protein localization [31]. Neurocalcin regulates sleep [57], but whether its myristoylation is involved in sleep regulation remains to be explored [58].

Farnesylation is an irreversible acylation which has primarily been studied for its tumorigenic properties on oncoproteins, like Ras [59]. Using a farnesyltransferase inhibitor to validate our farnesylation-specific antibody [33], we demonstrate that farnesylation levels change in the central brains of flies with sleep state (Supp. Figure 5C). We observed sexually dimorphic regulation by sleep, where males have elevated farnesylation levels when well rested, which was inversely correlated with changes in 15-carbon FA levels (Supp. Figure 5C). In females, 15-carbon FA levels increased with sleep need, which correlated with trending increases in farnesylation following sleep-deprivation (Supp. Figure 5D). This trending increase in farnesylation is associated with undefined, bright punctae (Supp. Figure 5E). While we do not know what these structures are, they could represent coalescing of toxic substances in small areas of the brain. Notably, farnesylation of RAS and ERK is elevated in individuals with Alzheimer’s disease (AD, and in a mouse AD model, memory impairments and other amyloid pathologies are suppressed with a knockout of farnesyltransferases [60]. Uch-L1, a protein associated with familial Parkinson’s disease, regulates mitochondrial dynamics and ER-mitochondria contacts in its farnesylated state, with conserved effects in flies [61]. Interestingly, Parkinson’s disease is highly associated with sleep dysfunction, and women have faster disease progression and mortality despite higher incidence in men [62].

While availability of FAs may directly regulate changes in protein acylation, if any, with sleep or wake conditions, FAs may also be activated via esterification using acetyl-CoA. We previously reported that acetylcarnitine increases with high sleep need [16]. Therefore, acetyl-CoA availability, acetylcarnitine production or acetylation of proteins could also serve to regulate acylation of proteins. One such enzyme that may mediate this is bubblegum, a fatty acyl-CoA synthetase, which prefers palmitic acid (FA 16:0) for thioesterification [63]. The DGs among our top hits in females with high sleep need often contain an esterified FA 16:0 chain, rather than its non-esterified (free) counterpart. *Bubblegum* has previously been implicated in sleep, with higher sleep rebound after sleep deprivation [12], and in regulating VLCFA levels [64], which also were associated with sleep need in this study (Figure 4D).

ULCFAs are a sub-type of VLCFAs identified by our brain-sorted lipidomics that are of particular interest. While there may be some regulation by sleep need in females, most striking differences in ULCFAs are observed by sex (Figure 4D). Neuronal ULCFAs were present only in females, while glial cells showed higher ULCFA levels in females than in males (Figure 4D). Although further research is needed to confirm the identification of these ULCFA species, a clear sex-specific pattern emerges. VLCFAs and ULCFAs are produced by the elongation of very long chain FAs (Elovl) family. There are even more *Elovl* genes in flies than in mammals, including some that are expressed in only one sex [65, 66]. Mutations in the Elovl family typically impact the brain, where VLCFAs and ULCFAs are found more than most other organs [28, 29]. FAs of these lengths must be broken down to shorter chain lengths by peroxisomes before undergoing mitochondrial beta-oxidation [28]. In flies and mammals, VLCFAs and ULCFAs become elevated with neurodegeneration [67], whereby they contribute to the activation of mouse astrocytes [68]. Thus, accumulation of VLCFA and ULCFAs could be pathogenic as they can induce neuroinflammation and shorten lifespan through glial accumulation. Indeed, reducing *Elovl1* function ameliorates these effects in a rodent model of multiple sclerosis (MS) [67]. While our study does not address FA transport between cell types, it is intriguing that female neurons accumulate VLCFAs with high sleep need, and this pattern is inverse to that of glia which have low VLCFAs with high sleep need (Figure 4D). Perhaps neuron-glia lipid coupling, including lipid transport and communication between these cells during sleep and wake, is also sexually dimorphic.

Finally, we report promotion of sleep by a class of lipids, monoacylglycerols (MGs), which are neutral storage lipids with a singular FA tail [30]. Feeding a multi-class MG lipase inhibitor, NF1819, has sex-specific effects on sleep, promoting sleep in females (Figure 6A and Supp. Figure 6A) [43]. NF1819 and other MG lipase inhibitors have been tested for treatment of MS [69, 70], an autoimmune disease that typically leads to severe loss of myelin [71]. Although fly neurons are not myelinated, myelin is made of multiple types of lipids, with the majority having VLCFA tails. Thus, myelin loss may be due to multiple mechanisms, including lipid metabolic processes conserved in fly glia [67]. Many people with MS have disordered sleep, especially insomnia, and their sleep disorders are often associated with brain region-specific demyelination [71], which can be prevented with MG lipase inhibitors [70]. Given that most people with MS are women [71], understanding sex-specific differences in glial lipid homeostasis and in glial MG signaling may be important for understanding this disease.

Studies of MG signaling have mainly focused on a sub-type of bioactive MGs called endocannabinoids, which are detected in *Drosophila*, though mammalian endocannabinoid receptors are not conserved [72]. However, non-endocannabinoid MG signaling was recently implicated in metabolic functions in the *Drosophila* brain [73], suggesting a role for various types of MGs. In our study, we show MGs can promote sleep in females by inhibiting MG lipases with NF1819 supplementation (Figure 6A). Although males do not increase sleep with NF1819 supplementation (Supp. Figure 6A), male glia have a greater DG signal than females, which does not shift noticeably from alert to sleep need conditions (Supp. Figure 4B). This is consistent with males having higher levels of *tuntid* expression than females [24]. Interestingly, the sleep-promoting *Fabp7* modulates brain endocannabinoid levels, particularly in female mice, indicating that female-specific sensitivity to MGs may be conserved [74]. Here, we find that MG accumulation promotes *Drosophila* sleep with glial knockdown of *tuntid* (Figure 6B-C). Flies upregulate *tuntid* expression, and thus DG synthesis, in response to starvation [24], which is consistent with starvation-associated sleep suppression [12]. We previously found that MGs protect against *Drosophila* seizures through TRP channels [75], which is the same family regulated by the *bishu-1* for larval cold avoidance [38]. We found similar seizure protection by supplementing NF1819 [42], suggesting regulation of glial TRP channels as a potential mechanism by which *tuntid* regulates sleep.

In summary, we uncover an important sleep-regulating role for the processing of neutral lipids in glia. The sex-specific regulation of lipids we report here has implications for understanding how metabolic needs are met differently by females and males, to support their specific behavioral patterns.

## Materials

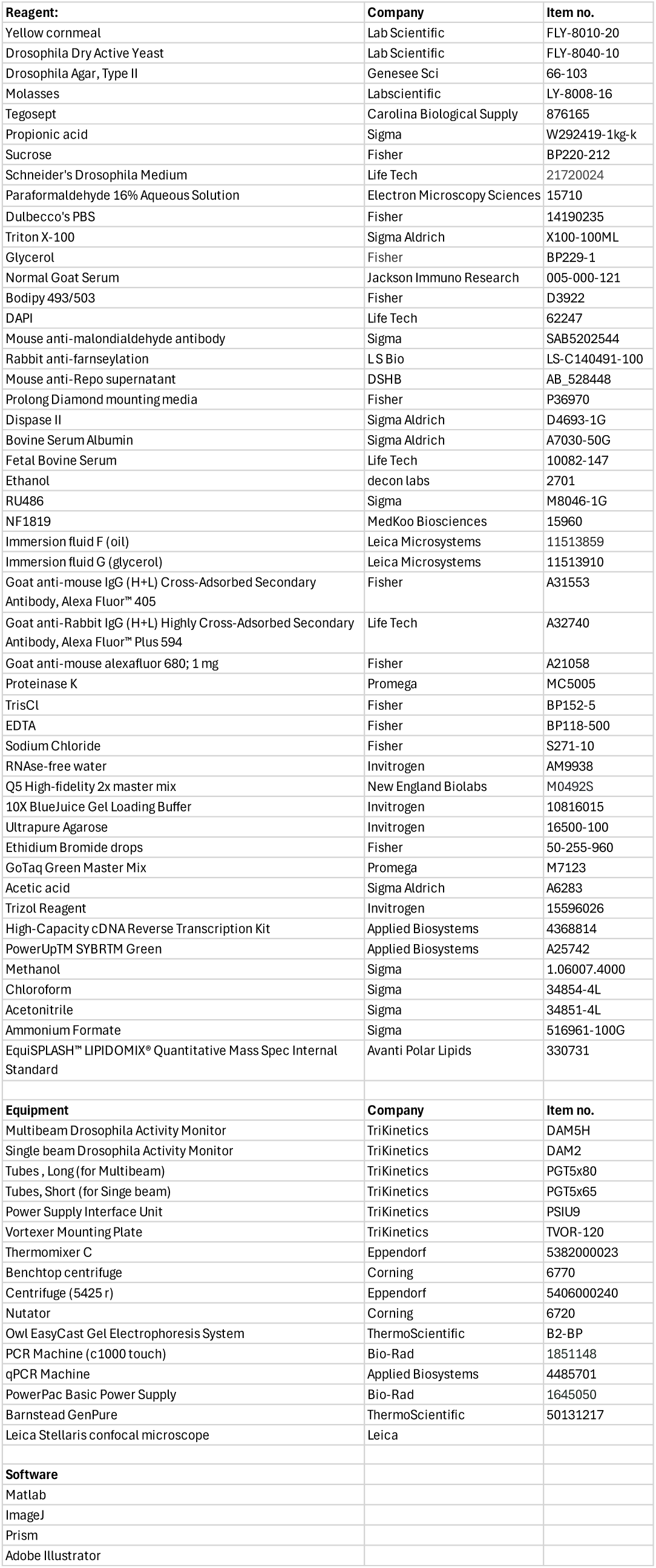

## Methods

### Fly Stocks

Fly stocks were obtained from Bloomington *Drosophila* Stock Center (BDSC), Vienna *Drosophila* Resource Center (VDRC), other labs, and lab stocks. The following fly lines were obtained from BDSC: *y^1^, lsd2^51^*/*FM7i* (#98125), y^1^ (#169), cortex-Gal4 (#45784, w^1118^;; PGMR54H02-Gal4attP2, cortex glia-specific Gal4 newly outcrossed in lab to iso^31^), w*;;P20XUAS-IVS-mCD8-GFPattP2 (#32194), inaE RNAi (#64885, y^1^sc*v^1^sev^21^; PTRiP.HMC05758attP40;+), P40 control (#36304, y^1^v^1^;PMsp300, attP40;+), and midgut Gal4 (#91368, w^1118^; Pmex1-Gal4.2.110-8, midgut-specific Gal4 newly outcrossed in lab to iso^31^). The following lines were obtained from VDRC: exo70 RNAi KK (#103717, y^1^w*; PKK101154VIE-260B;+), 30B control (#60103, KK UAS-GFP-RNAi control line for 30B landing site), bishu-1 RNAi KK (#107788, y^1^w*; PKK109140VIE-260B;+), bishu-1 RNAi GD (#7942, w^1118^;; PGD1854v7942), bishu-2 RNAi KK (#108495, y^1^w*;PKK112067VIE-260B;+), CG1941/tuntid RNAi GD (#3998, w^1118^, PGD1853v3998;;+), and rbo RNAi GD (#47751, w^1118^; PGD14013v47751;+). The following fly lines were obtained from other labs: w’;;bmm^1^/TM3 (Ronald Kuhnlein lab), Canton S (Leslie Griffith lab), and w* Canton S (wCS, Leslie Griffith lab). The following fly lines were obtained from Sehgal lab stocks: iso^31^, w’;;Repo-Gal4/TM6C (outcrossed to iso^31^), w’;; 20X-UAS-TTS-shi^TS^-p10/(TM6C) (outcrossed to iso^31^), w’;9137-Gal4;+ (outcrossed to iso^31^), w’;;Repo-Geneswitch (outcrossed to wCS), and w;;Repo-Gal4/TM6C (outcrossed to wCS). The following lines were newly generated by crosses for this manuscript, including control lines for mutant alleles made from siblings: y^1^, Lsd2^51^/FM7i;; Repo-Gal4/TM6C (crossed to iso^31^ and y^1^ once), y^1^;;Repo-Gal4/TM6C (crossed to iso^31^ and y^1^, Lsd2^51^ once), w’;; bmm^1^, Repo-Gal4/TM6C (mutant allele crossed to iso^31^ once), w’;;Repo-Gal4/TM6C (crossed to bmm^1^ once), w’;; bmm^1^, 20X-UAS-TTS-shi^TS^-p10/TM6C (mutant allele crossed to iso^31^ once), w’;; 20X-UAS-TTS-shi^TS^-p10/TM6C (crossed to bmm^1^ once), +;;Repo-Gal4/TM6C (newly crossed to Canton S), and +;;Nsyb-Gal4 (newly crossed to Canton S).

### Fly Rearing

Flies were raised and tested in a 12-hour light/12-hour dark cycle. Flies were mated and maintained at 25C and 50-70% humidity, except where otherwise noted. Sex varied by experiment, as noted. All sleep experiments display data from flies aged 6-12 days post-eclosion during dates averaged, while most other experiments use mated flies 6-9 days post-eclosion. Females and males were housed together before experiments to facilitate mating. Unless otherwise described, flies were fed the Sehgal Lab Standard Yeast-Molasses Diet, which we refer to as “nutrient-complete diet.” This diet includes the following: 64.7g/L corn meal; 27.1g/L dry yeast; 8g/L agar; 61.6mL/L molasses; 10.2mL/L 20% tegosept; 2.5mL/L propionic acid.

### Sample Preparation for Microscopy

Flies are raised on nutrient-complete food and transferred to fed fresh food within 24 hours of dissection. They were briefly anaesthetized with carbon dioxide before dissection in Schneider’s media. The brains are fixed immediately for 23-29 minutes in 4% paraformaldehyde in Phosphate Buffered Saline (PBS), then cleaned in 0.1% PBS-Triton-X 100 (PBST). The brains are washed in 0.1% PBST 2-3 times, totaling at least 30 minutes. 5% normal goat serum in 0.1% PBST (5% NSGT) is added to block the brains for one hour while nutating at 4C. After one hour, the blocking supernatant is removed and replaced with primary antibodies in 5% NGST. The brains remain nutating overnight in primaries before washing 3 times for 10 minutes each in 0.1% PBST. Secondary antibodies in 5% NGST are added after supernatant removal. The tubes are covered in foil before nutating overnight at 4C. The secondaries are washed 3 times for 10 minutes each in 0.1% PBST. The supernatant is replaced with 50% glycerol in 0.1% PBST until mounting within 2 weeks. The brains are mounted in ProLong Diamond Antifade mounting media, which cures at room temperature for at least 24 hours before long-term slide storage at 4C.

For experiments using mouse anti-malondialdehyde primary antibodies, the 1:100 antibody to 5% NGST concentration was added after NGST blocking. Far-red secondary antibody, AlexaFluor 680 was used to minimize detection of air tubules and cellular debris by autofluorescence at lower wavelengths or detection of wavelengths emitted by Bodipy 493/503 at a 1:200 secondary:5%NGST concentration.

All LD experiments were performed in conjunction with immunohistochemistry. For any LD experiments, 1000uL of 1:1000 Bodipy 493/503 in PBS was added to microcentrifuge tubes containing brains after washing secondary antibodies with 0.1% PBST twice. The brains rotated in Bodipy-PBS mix for 3-4 hours and then were immediately mounted in ProLong Diamond Antifade mounting media. Prior to mounting in ProLong Diamond Antifade, the brains are left in 80% glycerol (in PBS) for at least 30 minutes to dilute Bodipy carryover and mitigate brain desiccation during mounting. Slides were cured for at least 24 hours at room temperature before long-term storage at 4C.

For experiments using rabbit anti-farnesylation primary antibodies, 1:200 antibody:NGST concentration was added to the brains after NGST blocking. To set Z-limits based on brain cellular physiology, these brains were either co-stained with mouse anti-Repo primary antibodies (1:300; anti-mouse AlexaFluor 405 secondary antibodies, 1:1000) or stained with 1 ug/mL DAPI in 0.1% PBST during last 10 minute wash after secondary antibodies. anti-Rabbit AlexaFluor 594 secondary antibodies were used for all farnesylation experiments to minimize detection of air tubules and cellular debris by autofluorescence at lower wavelengths or detection of wavelengths emitted by DAPI and AlexaFluor 405. All imaging experiments were performed on a Leica Stellaris confocal microscope and all images were analyzed using ImageJ. 25-40 slices per brain were imaged, depending on experiment with a consistent number of slices per brain imaged per replicate. All LD experiments had 40 slices per brain and were imaged using a 40X oil objective. MDA experiments were imaged with 40X oil, 20X oil, or 20X glycerol objectives. Farnesylation experiments were imaged with 20X oil or 20X air objectives.

### LD Analysis

LDs were analyzed using methods and macros we published previously with minor adjustments to measure LD count, area, and average size [6]. Here, we drew regions of interest (ROIs) around central brains and performed analysis with macros adjusting settings for each replicate, as needed. Lipid trash was removed by macro exclusion only, and all images within a replicate were processed with the same macro settings. Within the macro, the triangle auto-threshold function was used on individual slices to generate a binary image stack marking lipid droplets for analysis. We report LD count and LD area as determined by the macro; average size is determined by dividing LD area by the LD count. For all LD experiments, metrics are normalized using the median value of the Gal4 control for its replicate. For experiments conducted at multiple time points, the ZT2 Gal4 control was used for normalization. For all representative images, projections of maximum intensity are displayed in green. Scale bars are included (50 or 100 microns, as noted). Some images include zoomed ROIs (enlarged 4.0x, Figure 3C) which were added using a publicly available ImageJ macro [76].

### Immunohistochemistry analysis

For all representative images, projections are summed and have brightness and contrast adjusted to have an identical range in pixel value. Images are false colored with magma (Fig 1F) or plasma (Supplementary Figure 6D-F) LUTs. Scale bars (50 or 100 microns, as noted) and color calibration bars are included. Some images include zoomed ROIs (enlarged 3.0x, Supplementary Figure 6E) which were added using a publicly available ImageJ macro [76]. Summed projections were analyzed determining a maximum threshold pixel value that could both distinguish the central brain from background as well as detect central brain regions across all conditions. The threshold was determined for each replicate by eye and by testing on multiple brains per condition before running. An ROI was drawn around each central brain, and average pixel intensity above the threshold is reported. Two ImageJ macros were written to perform the same analysis on and to output data from an entire replicate once the minimum threshold was determined and ROIs drawn. For MDA experiments using normalization, the ZT2 Gal4 control was used for normalization. For experiments quantifying farnesylation, the median value for ZT2 males was used for normalization.

### anti-Farnesylation Antibody Confirmation

Two farnesyltransferase inhibitors (FTI), tipifarnib and lonafarnib, were reconstituted in dimethyl sulfoxide (DMSO). 50 ug/mL of each was added to reheated nutrient-complete food for a FTI concentration of 100 ug/mL [33]. Upon solidification, adult, male Canton S (CS) flies (4-7 days post-eclosion) were fed freshly made FTI-supplemented diet for one week. Brains were dissected at ZT2 from +FTI and DMSO controls and prepared for confocal microscopy. Images were compared by qualitative analysis.

### Sleep Behavior

All sleep behavioral experiments use 14-35 flies per group per replicate, analyzing data from at least three replicates, except where noted. Flies were fed nutrient complete until being loaded into multibeam Drosophila Activity Monitors (DAM) tubes with 5% sucrose/2% agar food (referred to as sucrose or nutrient-depleted diet) 4-7 days post-eclosion. The first 36-48 hours allowed for adjustment to tube. Movement and position distributions were recorded from three days (days 2-4 post-loading), averaged, and used to calculate standard sleep and activity metrics. Dead flies and flies in tubes with mold or desiccation were excluded prior to analysis. Any channels with no detection of activity were excluded after analysis. DAMfilescan and previously reported custom MatLab scripts were used to calculate sleep metrics for all experiments [77].

If fed food containing RU486 (mifepristone), 100 uM RU486 is added to sucrose diet during the DAM tube preparation. The same volume of 200 proof ethanol is added for -RU486 controls. Flies were loaded at the same ages and analyzed in the same manner as other sleep experiments.

If fed nutrient-complete food in DAM tubes, flies were loaded 5-8 days post-eclosion and given 12-16 hours to adjust to food. Movement and position distributions were recorded from two days (days 1-2 post-loading) to mitigate larval movement being detected and analyzed as adult movement.

### NF181S Supplementation

Adult, CS flies 3-6 days post-eclosion are moved from a nutrient-complete diet to a nutrient-complete diet supplemented with either 500 ug/mL NF1819 (in ethanol) or an equivalent volume of ethanol [42]. Two days later, they are loaded into sucrose-based multibeam DAM tubes supplemented with the same concentration of NF1819 or ethanol as previously fed. The first 36-48 hours are discarded for adjustment to tube and diet, and the following three days of sleep are averaged and analyzed in the same manner as other sleep analyses described here using multibeam DAMs.

### Sleep Deprivation

Mechanical sleep deprivation consists of a random two seconds of mechanical shaking per twenty seconds. Flies undergo mechanical sleep deprivation for 10 hours beginning at ZT16 and ending at ZT2. Flies that underwent sleep deprivation for lipidomics or farnesylation experiments were deprived in vials. All flies are separated by sex 12-24 hours into fresh vials before mechanical sleep deprivation. Flies sleep deprived for behavioral assays are loaded into single-beam DAM tubes about 12-16 hours before baseline sleep recording begins. Analysis begins at the start of the next light cycle, recording one full light:dark cycle of baseline sleep, 10-hour sleep deprivation starting at ZT16 of the second light:dark cycle, and a final day of recording to analyze response to deprivation. This behavioral paradigm was used for neuronal and glial expression of mCD8-GFP sleep experiments.

### Sample Preparation for Cell Sorting

Samples for cell sorting were prepared using a modified protocol that was optimized for *Drosophila* brains [78]. Whole fly brains were dissected in cold Schneider’s medium at ZT2 (alert and sleep-deprived) or ZT14 (sleepy). 10 brains (Nsyb-Gal4>20XUAS-IVS-mCD8-GFP) for each neuronal condition and 60 brains (Repo-Gal4>20XUAS-IVS-mCD8-GFP) for each glial condition were dissected. Due to the large number of brains to be dissected for one sex in a short time, dissections were best facilitated by prior removal of one sex 12-24 hours before dissection. Brains were transferred to a microcentrifuge tube and spun down with a bench top centrifuge to remove Schneider’s media. 500ul of 1mg/mL Dispase II in Schneider’s medium was added to each tube. Digestion occurred on a thermoshaker, shaking at 1000 rpm for 15 minutes at 25C. The tubes were spun down with a benchtop centrifuge to remove dispase supernatant. The digested brains were rinsed three times with 200 ul of Schneider’s media, spinning down with benchtop centrifuge to remove tissue between washes. After final wash, 500 ul of 2% bovine serum albumin in phosphate buffered saline (PBS) was added to each tube holding brains. A 200ul pipette tip was coated with fetal bovine serum to prevent cells from sticking during physical cell separation. Pipet each sample up and down for 3 minutes, ensuring no large tissue chunks at end of time. Cells are transferred to a FBS-precoated sorting tube using a 40 micron filter. Tap the bottom of each sorting tube to mix after adding 1uL of 1mg/mL DAPI. Use fluorescence activated cell sorting (FACS) to enrich for DAPI-negative, GFP-positive cells. Sort cells into microcentrifuge tubes containing 10 ul of cold Schneider’s medium. After sorting, centrifuge cells at 300 x g for 3 minutes at -8C. Supernatant removed before long-term storage at -80C.

### Lipidomic Analysis via Flow Injection MRM Profiling

Flow injection-based multiple reaction monitoring (MRM) profiling was performed without chromatographic separation (i.e., direct infusion) using an Agilent 6495C triple quadrupole mass spectrometer (Agilent Technologies, Santa Clara, CA, USA) coupled to an Agilent 1290 Infinity II liquid chromatography system equipped with an Agilent Jet Stream (AJS) electrospray ionization (ESI) source. Approximately 3,000 lipid species were targeted across multiple lipid classes, including diacylglycerols (DG), triacylglycerols (TG), ceramides (Cer), cholesteryl esters (CE), acylcarnitines (CAR), free fatty acids (FA), glycerophosphocholines (PC), sphingomyelins (SM), glycerophosphoserines (PS), glycerophosphoinositols (PI), glycerophosphoethanolamines (PE), glycerophosphoglycerols (PG), and various lysoglycerophospholipid subclasses. Class-specific acquisition methods were employed, with DGs and TGs profiled using ammonium adducts and monitoring neutral losses corresponding to targeted fatty acyl chains. Lipids were extracted from biological samples using a modified Bligh and Dyer protocol [79], dried under vacuum using a SpeedVac, and reconstituted in 200 µL of 3:1 methanol:chloroform.

Prior to analysis, reconstituted lipid extracts were diluted 2-fold in 70:30 methanol:acetonitrile containing 10 mM ammonium formate and 0.1 µg/mL EquiSPLASH™ LIPIDOMIX® Quantitative Mass Spec Internal Standard (Avanti Polar Lipids, Alabaster, AL, USA), a stable-isotope-labeled mixture of deuterated lipid species spanning major lipid classes, designed for normalization and relative quantitation in lipidomics workflows. As a quality control (QC) measure, solvent blanks containing EquiSPLASH without lipid extracts were analyzed periodically throughout the run to monitor instrument performance and signal stability over time.

Lipid annotation, data parsing, statistical analysis, and visualization were carried out using the Comprehensive Lipidomic Automation Workflow (CLAW) MRM Platform [80], with all associated data files available on GitHub: https://github.com/cbeveri/drosophilia_brain_cell-sorted_lipidomics_across_sleep-wake.

### Genotyping

In order to recombine transgenic lines with amorphic alleles, we genotyped whole flies or wings of flies for *bmm^1^*, *Lsd2^51^*, and Repo-Gal4 alleles. Genomic DNA was extracted by adding 0.2 mg/ml proteinase K to squishing buffer. Squishing buffer also includes 10mM TrisCl (pH 8.0), 1mM EDTA, and 25mM NaCl. 10ul (wing genotyping) or 50 uL (whole-fly genotyping) of squishing buffer was added to a tube for DNA extraction. Wings were added and pressed down into solution and not further disrupted. Extraction from whole flies included squishing the fly with buffer-filled pipet tip in tube before expelling remainder of squishing buffer. Then the solutions were incubated at 37C for 1 hour followed by 95C for 5 mins. Solution was spun down and kept at -20C for long-term storage.

Q5 High-Fidelity 2X Master Mix was used to genotype *bmm* and *Lsd2*. The reaction included 12.5 ul of master mix, 1.25 ul forward primer, 1.25 ul reverse primer, 1.5-2.5 ul DNA, and nuclease free water to total 25ul per sample. *bmm* polymerase chain reaction (PCR) is as follows: 1) denaturation at 98C for 90 seconds, 2) denaturation at 98C for 10 seconds, 3) annealing at 62C for 20 seconds, 4) extend at 72C for 4.5 mins, 5) repeat steps 2-4 39 times, 6) final extension at 72C for 2 minutes. Lsd2 PCR is as follows: 1) denaturation at 98C for 30 seconds, 2) denaturation at 98C for 10 seconds, 3) annealing at 62C for 20 seconds, 4) extend at 72C for 1.5 mins, 5) repeat steps 2-4 34 times, 6) final extension at 72C for 2 minutes. Gel electrophoresis was conducted using 0.7% agarose gel made with 1xTAE buffer and ethidium bromide; 10X Blue Juice as added as loading dye to reactions and best band separation occurred after a couple hours at 70-95 Volts.

GoTaq Green Master Mix was used to genotype for presence of Gal4. The reaction included 12.5 ul of GoTaq Green Master Mix, 1 ul of forward primer, 1 ul of reverse primer, 2.5 ul of DNA and 8 ul of nuclease-free water. Gal4 reaction is as follows: 1) denaturation at 95C for 2 minutes, 2) denaturation at 95C for 30 seconds, 3) annealing at 61C for 30 seconds, 4) extend at 72C for 45 seconds, 5) repeat steps 2-4 34 times, 6) final extension at 72C for 5 minutes. Gel electrophoresis was conducted using 1% agarose gel made with 1xTAE buffer and ethidium bromide; gels were run for 30 minutes at 120V.

### qPCR

Mated, female flies (4-5 days post-eclosion) were flash frozen on dry ice between ZT2-3. Their guts were dissected in Schneider’s media and pooled into groups of 20 guts per genotype per replicate. To confirm RNAi knockdown efficiency, RNA extraction was performed on guts samples. RNA was extracted using TRIzolTM Reagent and reverse transcribed to 1ug of cDNA using High-Capacity cDNA reverse Transcription kit.

Quantitative PCR (qPCR) analysis was performed using PowerUpTM SYBRTM Green on QuantStudioTM 7 Flex Real-Time PCR machine. Relative gene expression was calculated using the change in cycling threshold (△△Ct) method normalizing to tubulin.

### Primers

Primers for genotyping and qPCR include:

*bmm* Forward: GCAATGGCCTGAACAATAAAC

*bmm* Reverse: GTCGTGTGAGGGTGACTAATA

*Lsd2* Forward: ACACACGCATACCCTAGATTC

*Lsd2* Reverse: GTAACTCTACCGTCAACACTCAC

*Gal4* Forward: AAAGACGCCGAATTTGGTGGTC

*Gal4* Reverse: TGTGATAACAGAAACGGCGCAC

*tuntid* Forward: CGTGGTTCAGAGCGATCAA

*tuntid* Reverse: CATCTTCTCCAGGTCGTCAATAA

*alpha-tubulin 84B* Forward: CGTCTGGACCACAAGTTCGA

*alpha-tubulin 84B* Reverse: CCTCCATACCCTCACCAACGT

### Statistics

For all experiments, except those related to lipidomics analysis, Prism was used to conduct all statistical testing and produce plots. All quantifications are displayed with box plots where the box surrounds points within the interquartile range, a line within the box representing the median, and bars extending to total range of data points, except where noted. Higher n’s allowed reliable Shapiro-Wilkes pre-testing of all groups’ normality. This consistently showed that non-parametric testing was appropriate for sleep behavioral replicates, using Mann-Whitney test or Kruskal-Wallis test that is uncorrected because one or more control groups is compared to one experimental group. For imaging experiments, data that are normal use Welch’s t tests, comparing one or more groups to a single control. Non-parametric data use Kruskal-Wallis test that is uncorrected because one or more control groups is compared to one experimental group. For experiments with multiple variables, two-way ANOVA is used and p-values for the within-ZT or within-sex Dunnett’s post hoc test comparing one control with one experimental group is displayed. For qPCR data, a one-way ANOVA is used with Fisher’s post-hoc test with two control groups compared to one experimental group. Lipidomics data were analyzed using the CLAW MRM platform, with log fold-change and significance testing performed via the edgeR generalized linear model (GLM) [80].

## Contributions

ESP, PRH, and AS conceived the study. ESP, CB, PRH, VAK, and CER developed and validated methodology, as needed. ESP performed most experiments with assistance from VAK, SLK, ZY, ENA, KNL, and PAP. ESP analyzed data from most experiments with assistance from PRH and VAK. PRH and VAK wrote macros used for imaging analysis with minor editing by ESP. PCC performed and analyzed all qPCR experiments. All brains were dissected by ESP, SLK, and ZY, and processed by ESP and VAK for FACS sorting and lipidomics analysis. SGN advised on FACS-isolation from brain cells. CER performed mass spectrometry-based lipidomics experiments. CB wrote lipidomics analysis code and performed all lipidomics analysis. ESP, CB, CER, SI, KAW, and GC advised on lipidomics analysis and generation of related figures. ESP generated most non-lipidomics figures with additional figures generated by PCC and SGN. CB generated most lipidomics figures with assistance from SI and minor editing by ESP. ESP designed all graphics in Adobe Illustrator. ESP wrote the manuscript. CB, PRH, VAK, CER, PCC, SGN, KAJ, GC, and AS revised the manuscript. All authors contributed to data interpretation and reviewed the manuscript.

## Acknowledgements

This work was supported by the National Science Foundation’s Graduate Research Fellowship Program (ESP), the Bawd Foundation’s David Elliman Research Internship (VAK), the Arnold O. Beckman Postdoctoral Fellowship program (CER), the AnalytiXIN Fellowship Award (GC and CER), and Howard Hughes Medical Institute (AS). GC is the James Tarpo Jr. and Margaret Tarpo Professor at Purdue University. The work was also supported, in part, by NIH National Center for Advancing Translational Sciences U18TR004146 award, ASPIRE Challenge and Reduction-to-Practice awards, and Defense Threat Reduction Agency (DTRA) AIMS-HITS contract award MCDC2202-003 to G.C. The Purdue University Center for Cancer Research funded by NIH grant P30 CA023168 is also acknowledged. The authors would like to thank Agilent Technologies Inc. for the donation of the Triple Quadrupole LC/MS instrument to Chopra Laboratory. Data for this manuscript were generated in the Penn Cytomics and Cell Sorting Shared Resource Laboratory at the University of Pennsylvania (RRID:SCR 022376). Penn Cytomics is partially supported by the Abramson Cancer Center NCI Grant (P30 016520). We thank Cynthia Hsu for writing and teaching use of Matlab scripts for sleep behavioral analysis, China Burns and Sara Bernardez Noya for sharing their Drosophila brain-cell sorting protocols to assist in protocol optimization, and E. Evonne Jean for providing immunoassay advice. Lastly, we thank Sehgal Lab members, Chopra Lab members as well as David Raizen, Joe Baur, Erika Holzbaur, Daniel Rader, John Vaughen, and Lydia Grmai for their discussions and advice. The content is solely the responsibility of the authors and does not necessarily represent the official views of the National Institutes of Health.

