## Supplemental Figures and Legends for "Neutral lipid processing in glia is sexually dimorphic and promotes sleep through diacylglycerol catabolism"

**Supplemental Figure 1: LD storage nor malondialdehyde levels drive sleep associated with blocking transport in glial subtypes.**

Flies with a cortex glia-specific knockdown of exocytosis (iso/$y^1$w*; exo70 RNAi/+; cortex-Gal4/+) were assayed for MDA and LDs for comparison to the Gal4 (iso/$y^1$w*;30B/+; cortex-Gal4/+) and UAS (iso/$y^1$w*;exo70 RNAi/+) controls. A) Brain malondialdehyde levels, B) normalized LD count, and C) normalized LD area show no significant changes compared to both controls. Flies with a blood brain barrier (BBB) glia-specific transport block (9-137-Gal4>20X-UAS-TTS-$shi^{TS}$ -p10) were assayed for sleep, LDs, and MDA for comparison to Gal4 (9-137-Gal4>iso) and UAS (iso>20X-UAS-TTS-$shi^{TS}$-p10) controls. D) Sleep trace averaging three baseline sleep days for female flies fed a sucrose diet at room temperature in multibeam monitors. Normalized lipid droplet E) count, F) area, and G) size as well as H) normalized MDA levels are shown for an individual fly's central brain at ZT2 (yellow background) and ZT14 (grey background). A has n = 1 replicate with data points representing individual brains; Welch's t tests. B and C have n = 3 replicates; Kruskal-Wallis with Dunn's post hoc test, uncorrected. Its data points are normalized to Gal4 median value from its respective replicate, and all individual brains from all replicates are displayed. The values used for normalization for each replicate are the following: Count (B): 36212, 59887, 36662; Area (C): 1168918, 3500992, 1824261. D has n = 1 replicate; Kruskal-Wallis with Dunn's post hoc test, uncorrected. For E-H, all data points are normalized to Gal4 ZT2 median value from its respective replicate, and all individual brains from all replicates are displayed as data points. E-G have n = 4 replicates and H has n = 3 replicates; two-way ANOVA with within-ZT Dunnett's post hoc test. The values used for normalization for each replicate are the following: Count (E): 14177, 15568.5, 12959, 19308.5; Area (F): 409440, 548036.5, 931427.5, 1678958.5; Size (G): 31.2, 36.5, 72.5, 80.8; and MDA (H): 1449.8, 621.7, 248.5. For all data shown *$p \leq 0.05$, ** $p \leq 0.01$, *** $p \leq 0.001$, and **** $p \leq 0.0001$, while $p \geq 0.05$ is not significant (ns).

**Supplemental Figure 2: Sexually dimorphic sleep patterns of *bmm* mutants are observed on multiple diets**

Sleep trace and total sleep for *$bmm^1$* mutants (w'/w(iso);;$bmm^1$, Repo-Gal4/$bmm^1$) and controls (w'/w(iso);;Repo-Gal4/+) fed a nutrient-complete diet in single beam monitors. 2. A) No change in sleep is observed in females and B) males have decreased sleep. A has n = 2 replicates; Mann-Whitney test (two-tailed). B has n = 3 replicates; Mann-Whitney test (two-tailed). All figures display results from one representative replicate with data points representing individual flies. For all data shown *** $p \leq 0.001$, and $p \geq 0.05$ is not significant (ns).

**Supplemental Figure 3: Sleep behavior and gating conditions or flies used in sorted brain cell-lipidomics**

Baseline and homeostatic response to 10-hour mechanical sleep deprivation shown for A) females and B) males expressing mCD8-GFP in neurons [+/(w' or Y);;Nsyb-Gal4/20X-UAS-mCD8-GFP] and glia [+/(w' or Y);;Repo-Gal4/20X-UAS-mCD8-GFP]. C) Gating scheme for sorting approximately 100,000 DAPI-negative, GFP-positive neurons (above) and glia (below) from other brain cells. Representative neuronal and glial (sleep-deprived) gating schemes from the second female replicate are shown. D) Percentages of DAPI-negative, GFP-positive cells collected for each condition out of total cells sorted for four female (above) and male (below) replicates. Each color in each figure represents a different replicate collection. GSD: sleep-deprived glia, GSlpy: sleepy glia (ZT14), GAlrt: alert glia (ZT2), NSD: sleep-deprived neuron, NSlpy: sleepy neuron (ZT14) and NAlrt: alert neuron (ZT2). A-B have n = 3 replicates. D has n = 4 replicates per sex, totaling 8 collections. E) Principal component analysis showing sleep and sex conditions for all neuronal (left) and all glial (right) samples. Sleep conditions are indicated by color: Alert (blue), Sleep Deprived (SD, coral), and Sleepy (green). Sex is indicated by shape: Female (circle) and Male (triangle).

**Supplemental Figure 4: Sleep and sex interact to regulate lipid homeostasis, including levels of neutral storage lipids**

A) LogFC heatmap showing differentially expressed lipid species between conditions with high sleep need and alert flies. Comparisons with sleep-deprived (striped) and sleepy (grey) flies are outlined in green for males and pink for females. Brown represents a LogFC of $\geq$1 in sleep-deprived or sleepy compared to alert for each condition, and teal represents a LogFC of $\leq$-1. Non-significant lipids are white. B) Average DG and TG average ion intensities are visualized by sex, with males on the left and females on the right, and neurons displayed above glia. Increased DG levels are apparent in higher sleep need conditions (sleep-deprived and sleepy) compared to alert conditions in both male and female samples across both neuronal and glial cell types.

**Supplemental Figure 5: Farnesylation and other acylation-related fatty acids are regulated by both sleep and sex**

A) Relative intensities of fatty acid species with 2, 14, 15, and 16 carbons across all conditions. The lipid species of each condition with the highest intensity is set to 100%. Neuronal samples are displayed above glial samples for female sleep-deprived (coral), male sleep-deprived (gold), female sleepy (green), male sleepy (turquoise), female alert (periwinkle), and male alert (pink) conditions. B) Control flies were treated with DMSO, while +farnesyltransferase inhibitors (+FTI) were supplemented both tipifarnib and lonafarnib. Summed projections of confocal imaging of anti-farnesylation antibody staining with 100 micron scale bar. Images are false colored with plasma LUT. n = 3 control and n = 5 (+FTI) brains. C) Quantification of averaged anti-farnesylation antibody signal above a minimum threshold from central brains of wildtype Canton S flies that were dissected at ZT2 (alert, yellow background), ZT14 (sleepy, grey background), and ZT2 following 10 hour sleep deprivation (SD, striped background). n = 3 replicates D) Summed projections with of representative male brains from confocal imaging of anti-farnesylation antibody staining with 100 micron scale bar. Images are false colored with plasma LUT. E) Maximum projections of representative sleep-deprived females with 100 micron scale bar. SD females have bright punctae which were almost exclusively observed in this group. Images are false colored with plasma LUT. Arrows point to bright punctae with a small green rectangle is drawn around an ROI with multiple punctae. The image is enlarged 3.0 times and found in the red rectangle at the top right corner. For C), all data points are normalized to the male ZT2 median value from its respective replicate. n = 3 replicates, and data points represent individual brains from all replicates. The values used for normalization for each replicate are the following with thresholding value in parentheses: 1395 (400), 518.3 (110), 673.5 (225). Two-way analysis of variance (ANOVA) with within-sex Dunnett's post hoc test. For all data shown *$p \leq 0.05$, ** $p \leq 0.01$, *** $p \leq 0.001$, and **** $p \leq 0.0001$, while trending values $0.05 \leq p \leq 0.1$ are displayed, and $p < 0.1$ is not significant (ns).

**Supplemental Figure 6: Manipulating MG metabolism via multiple MGATs and diets alters sleep.**

Averaged multibeam sleep traces and total sleep on sucrose food. A) 500 ug/mL NF1819 supplementation has no or opposite effects on Canton S male sleep compared to ethanol control diet. B) Fold change of *tuntid/CG1941* mRNA expression is reduced comparing female guts with midgut-specific knockdown [purple; w', tuntid RNAi/$w^{1118}$ (iso);mex1-Gal4/+;+] to Gal4 controls [light grey; $w^{1118}$(iso); mex1-Gal4/+;+] and UAS controls [dark grey; w', tuntid RNAi/$w^{1118}$(iso);;+]. C) and D) Pan-glial knockdown of *bishu-1/CG1942* [$w^{1118}$(iso)/$y^1$w';;Repo-Gal4/bishu-1 RNAi KK construct] compared to Gal4 [$w^{1118}$ (iso)/$y^1$w';;Repo-Gal4/30B insertion control] and UAS [$w^{1118}$(iso)/$y^1$w';;bishu-1 RNAi KK construct/+] controls. Trending or significantly increased sleep is observed in C) female and D) male replicates. E-H) Pan-glial knockdown of *bishu-1* [$w^{1118}$(iso)/ w';;Repo-Gal4/bishu-1 RNAi GD construct] compared to Gal4 [w'(iso)/w';;Repo-Gal4/+] and UAS (w'(iso)/$y^1$w;;bishu-1 RNAi KK) controls. E) Females have no change in sleep and F) males inconsistently show reduced or no change in sleep when fed a sucrose-based, nutrient-depleted diet. G) Females have increased sleep and H) males inconsistently show reduced or no change in sleep when fed a nutrient-complete diet in single beam DAM monitors. For sleep-related figures, results from one representative replicate are displayed with data points representing individual flies. For the qPCR-related figure, data points represent *tuntid* mRNA levels for each replicate normalized to *tubulin* levels in the same sample and then to the UAS control, which was set to a value of 1. Error bars represent the standard error of the mean. A has n = 3 replicates; Mann-Whitney test (two-tailed). B has n = 3 replicates; one-way with Fisher's post-hoc test, uncorrected. C-D have n = 3 replicates; Kruskal-Wallis test with Dunn's post hoc test, uncorrected. E-F have n = 2 replicates; Kruskal-Wallis test with Dunn's post hoc test, uncorrected. G-H have n = 3 replicates; Kruskal-Wallis test with Dunn's post hoc test, uncorrected. For all data shown *$p \leq 0.05$, ** $p \leq 0.01$, *** $p \leq 0.001$, and **** $p \leq 0.0001$, $p \geq 0.05$ is not significant (ns).

**Supplemental Table 1**

Table S1. Lipidomics profiling of wild-type neurons and glia flies with increased sleep need (sleepy and sleep-deprived) versus alert flies of the same sex and cell type, related to Figure 5 and Table 1. Positive LogFC correspond with upregulation of the lipid in conditions of high sleep need.

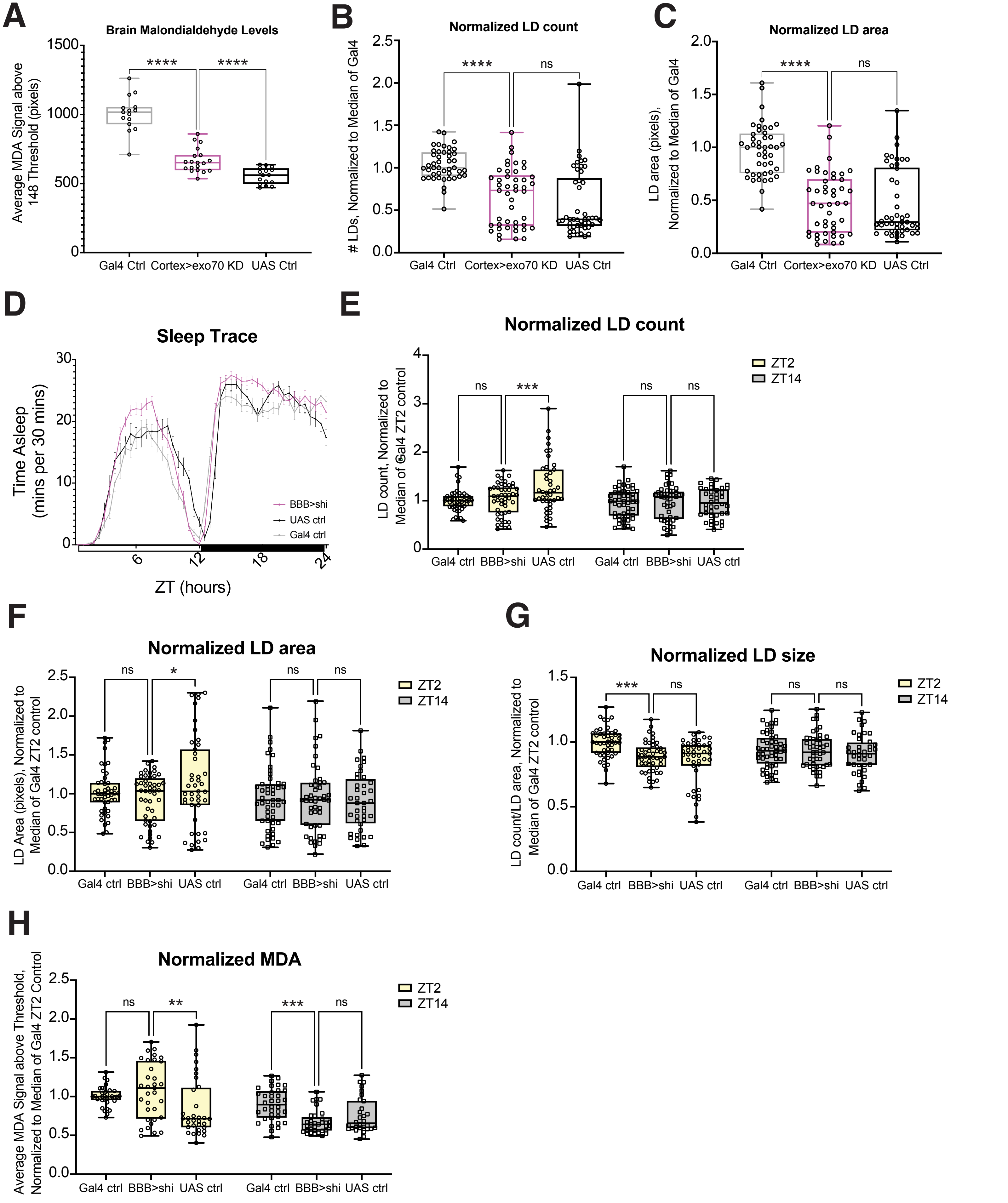


**Supplemental Figure 1: LD storage nor malondialdehyde levels drive sleep associated with blocking transport in glial subtypes.**

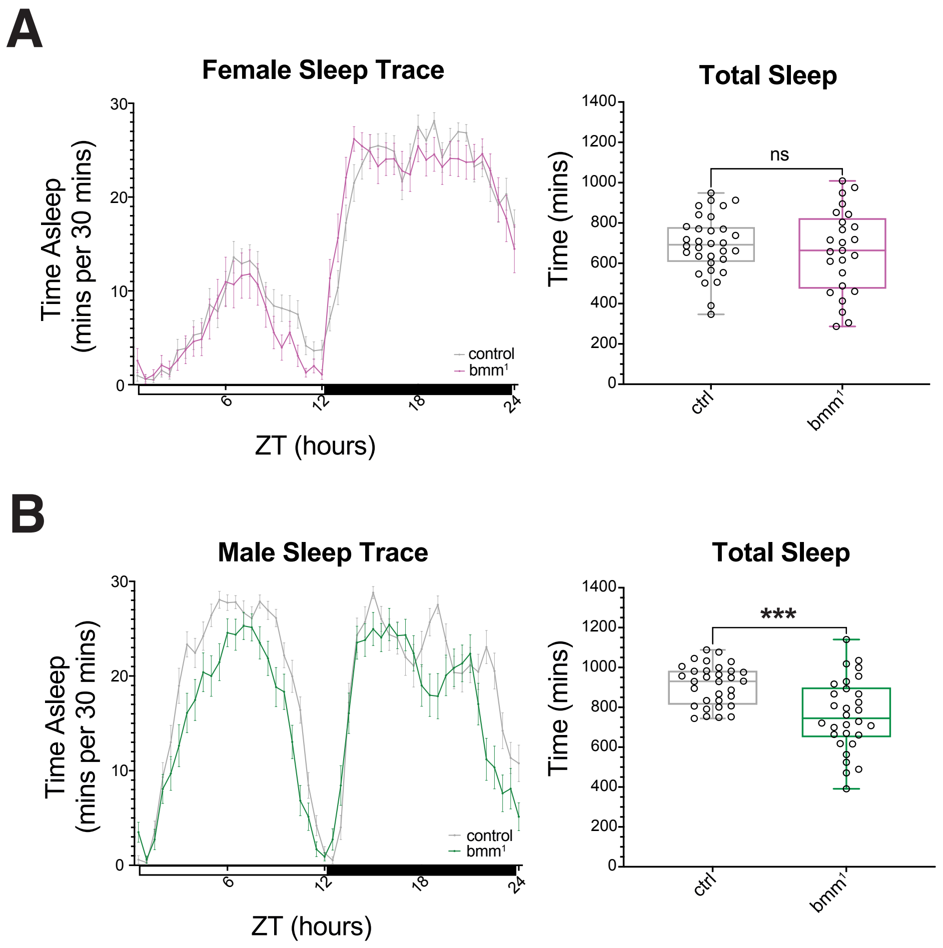


**Supplemental Figure 2: Sexually dimorphic sleep patterns of *bmm* mutants are observed on multiple diets**

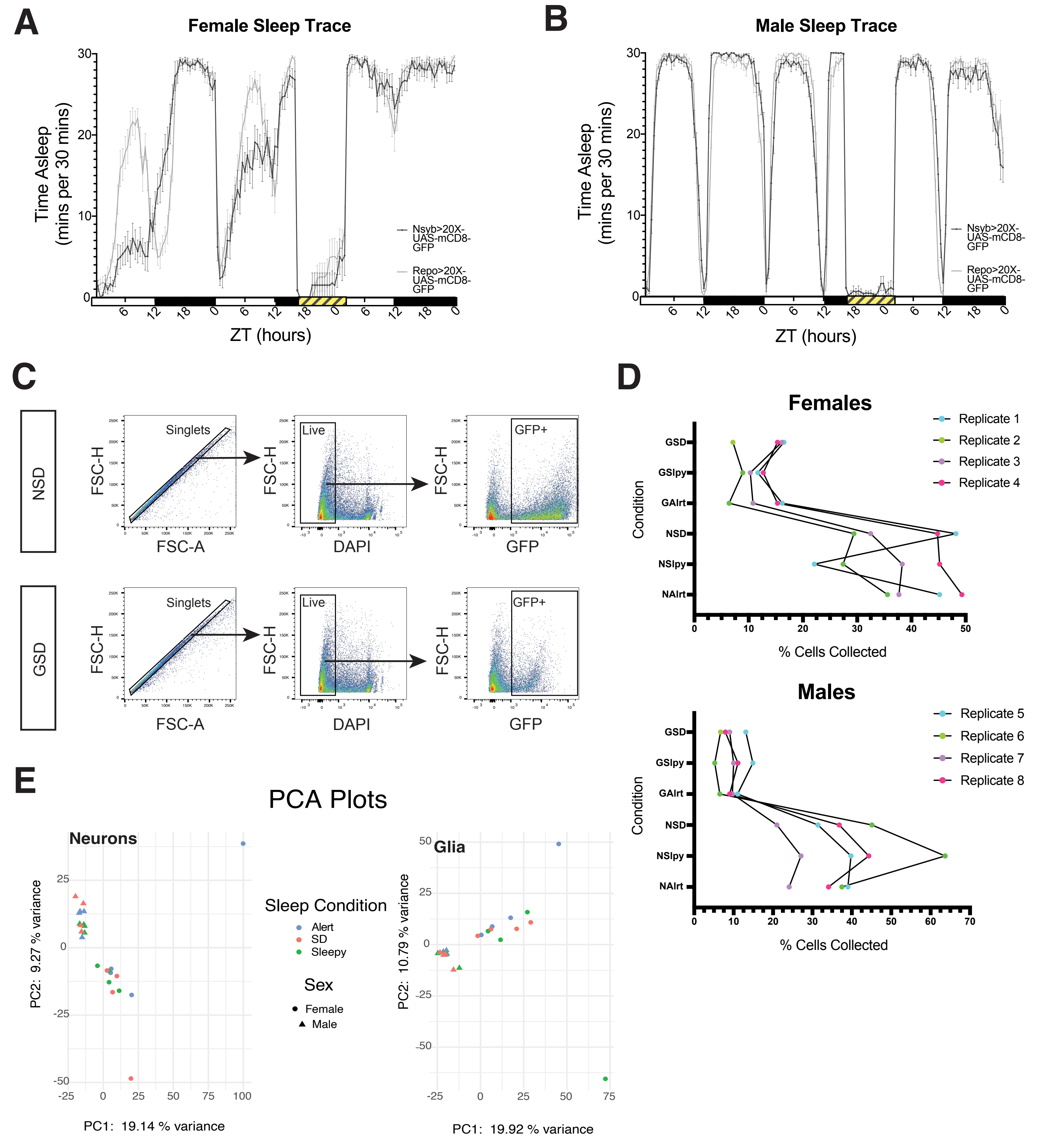


**Supplemental Figure 3: Sleep behavior and gating conditions for flies used in sorted brain cell-lipidomics**

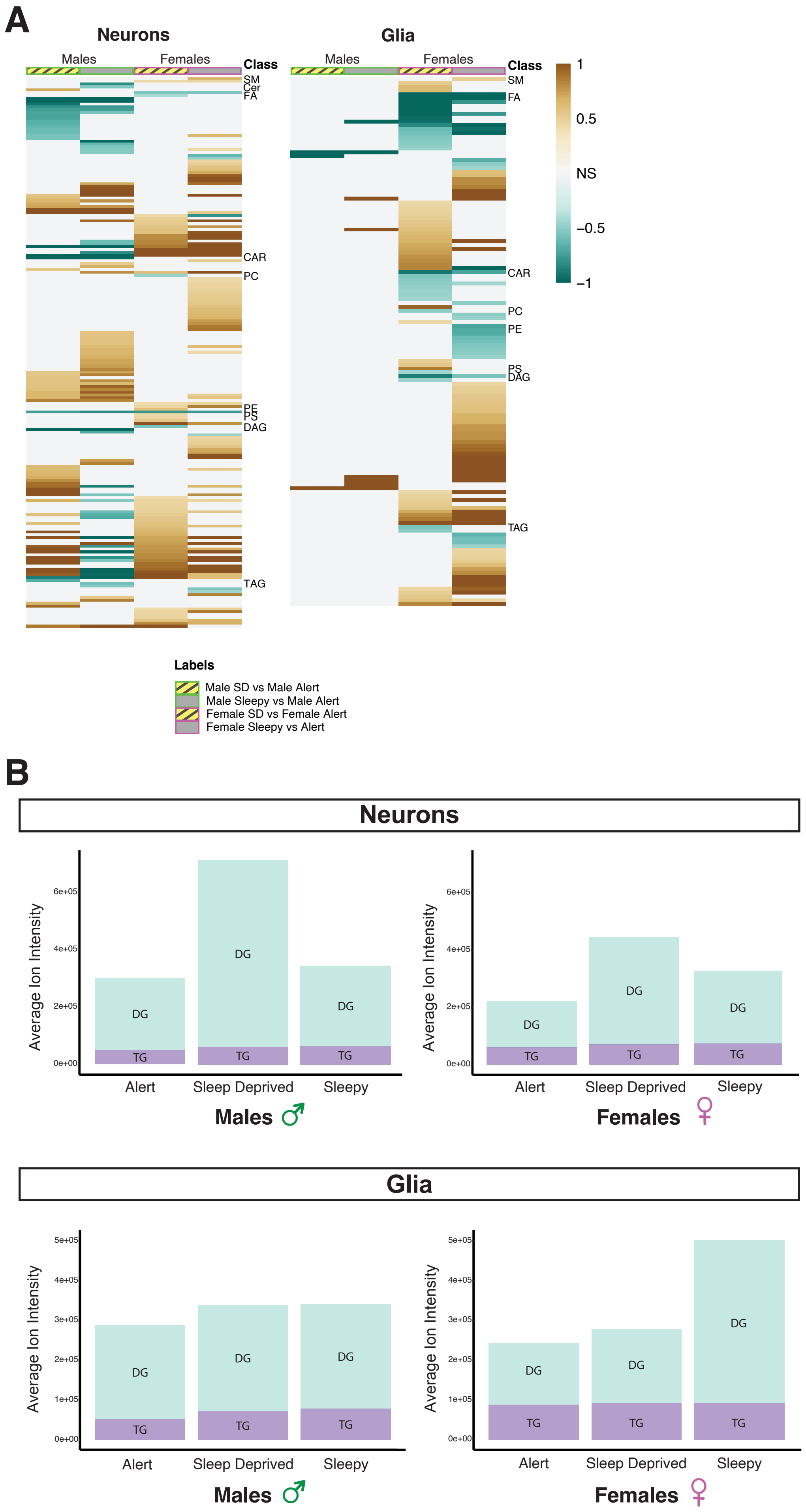


**Supplemental Figure 4: Sleep and sex interact to regulate lipid homeostasis, including levels of neutral storage lipids**

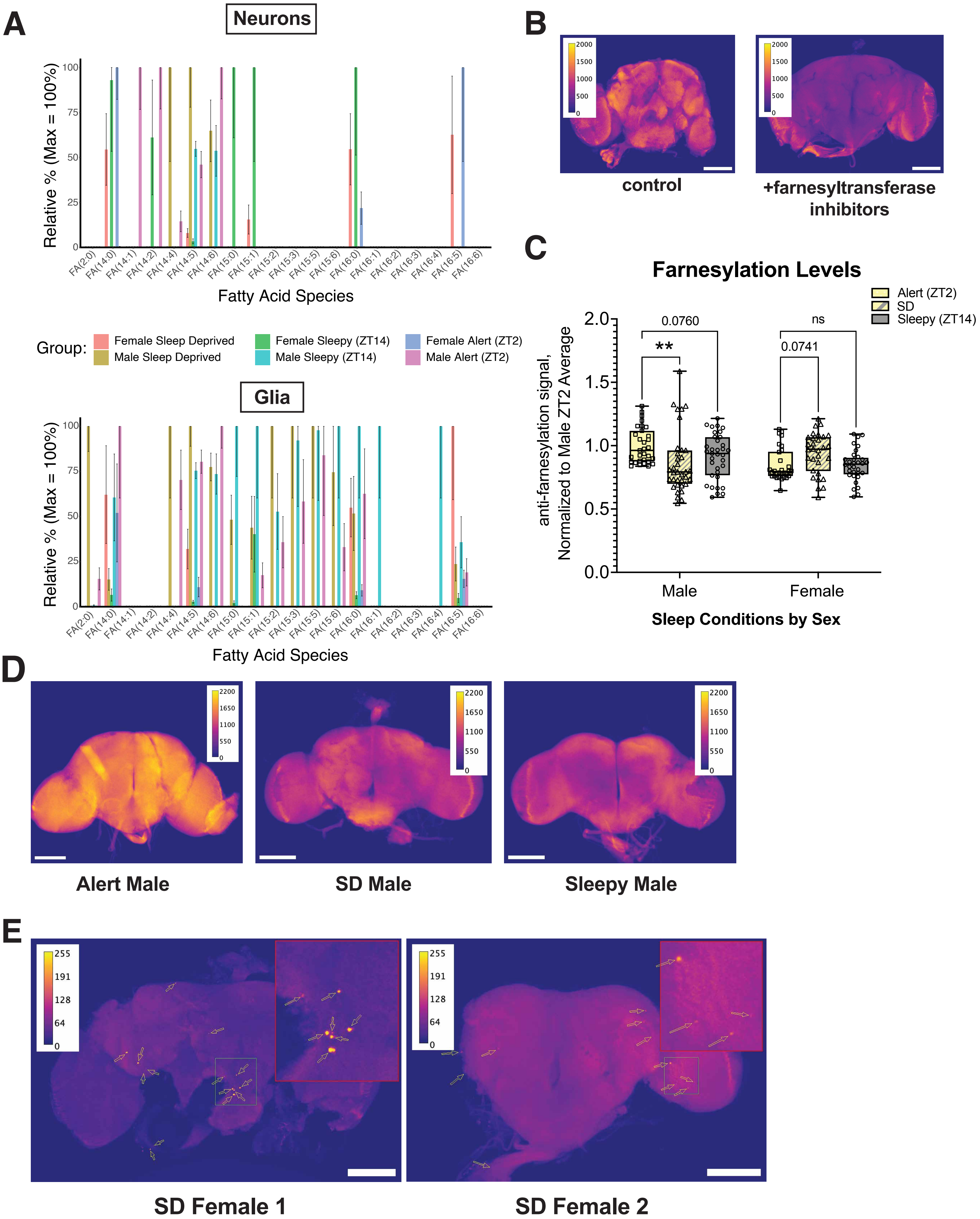


**Supplemental Figure 5: Farnesylation and other acylation-related fatty acids are regulated by both sleep and sex**

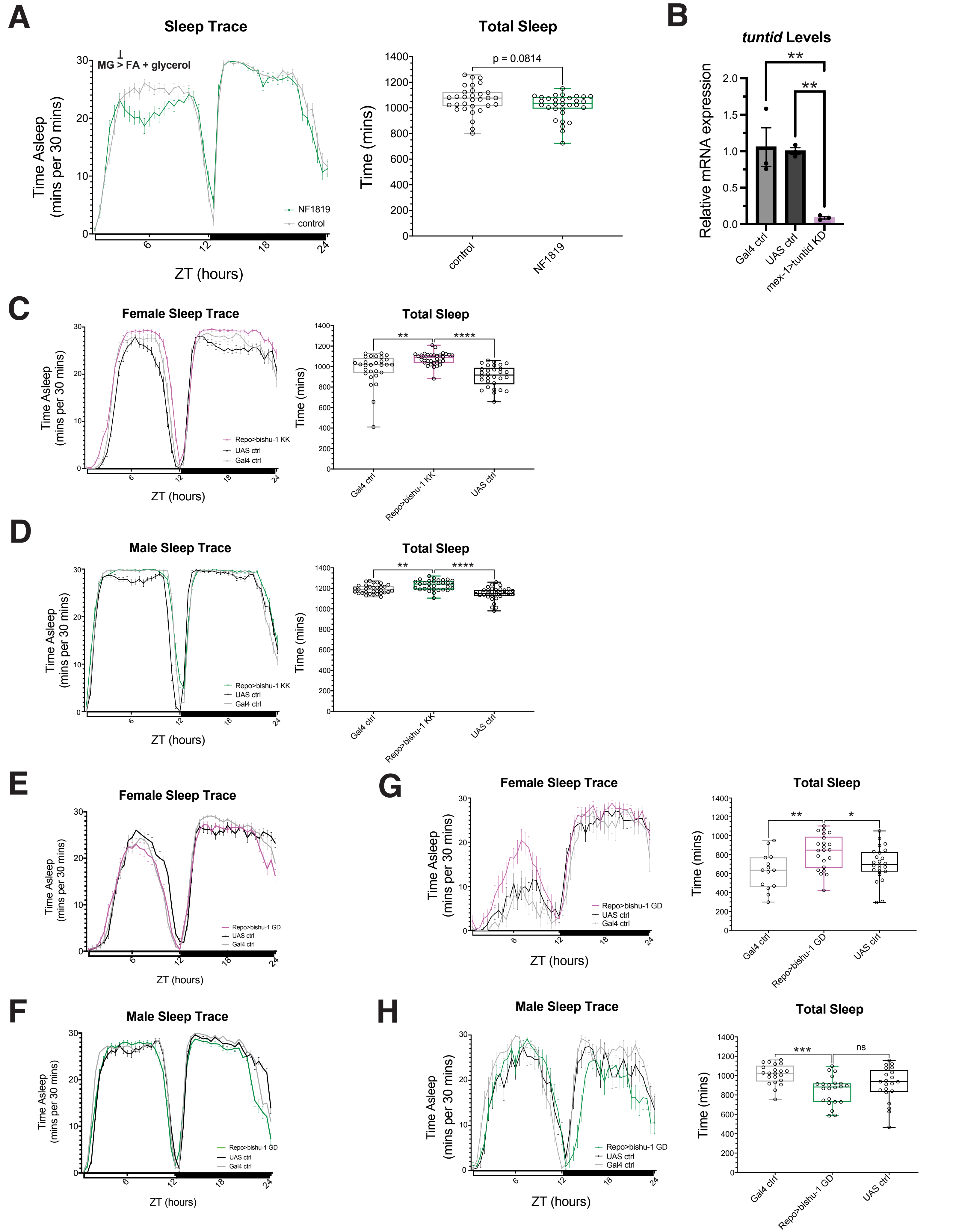


**Supp. Figure 6: Manipulating MG metabolism via multiple MGATs and diets alters sleep**
